# Cerebral organoid proteomics reveals signatures of dysregulated cortical development associated with human trisomy 21

**DOI:** 10.1101/315317

**Authors:** Tristan D. McClure-Begley, Christopher C. Ebmeier, Kerri E. Ball, Jeremy R. Jacobsen, Igor Kogut, Ganna Bilousova, Michael W. Klymkowsky, William M. Old

## Abstract

Human trisomy 21 (Down syndrome) is the most common genetic cause of intellectual disability, and is associated with complex perturbations in protein expression during development. Brain region-specific alterations in neuronal density and composition originate prenatally in trisomy 21 individuals, and are presumed to underlie the intellectual disability and early onset neurodegeneration that characterizes Down syndrome. However, the mechanisms by which chromosome 21 aneuploidy drives alterations in the central nervous system are not well understood, particularly in brain regions that are uniquely human and thus inaccessible to established animal models. Cerebral organoids are pluripotent stem cell derived models of prenatal brain development that have been used to deepen our understanding of the atypical processes associated with human neurobiological disorders, and thus provide a promising avenue to explore the molecular basis for neurodevelopmental alterations in trisomy 21. Here, we employ high-resolution label-free mass spectrometry to map proteomic changes over the course of trisomy 21 cerebral organoid development, and evaluate the proteomic alterations in response to treatment with harmine, a small molecule inhibitor of the chromosome 21 encoded protein kinase DYRK1A. Our results reveal trisomy 21 specific dysregulation of networks associated with neurogenesis, axon guidance and extracellular matrix remodeling. We find significant overlap of these networks show significant overlap with previously identified dysregulated gene expression modules identified in trisomy 21 fetal brain tissue. We show that harmine leads to partial normalization of key regulators of cortical development, including WNT7A and the transcription factors TBR1, BCL11A, and POU3F2, pointing to a causative role for DYRK1A over-expression in neurodevelopmental effects of human trisomy 21.

## INTRODUCTION

Trisomy of chromosome 21 (HSA21), the cause of Down syndrome (DS), is the most common viable human aneuploidy with an estimated prevalence of ~8 in 10,000 live births (1). DS is characterized by a complex spectrum of clinical manifestations that includes intellectual disability, cardiovascular and craniofacial abnormalities, early onset dementia with Alzheimer’s neuropathology (2), increased risk of childhood leukemias, and reduced risk of solid tumors (3). Among the most consistent neuroanatomical phenotypes in DS is a reduction in the surface area of specific regions of the cerebral cortex (4). These alterations have been attributed to disruption in neurogenesis and synaptogenesis programs during fetal brain development (5-9). Hypocellularity in fetal (6, 10) and adult DS neocortex (11), and defective lamination in the developing neocortex (12) suggest that atypical neuroanatomical features in adult DS brain originate early in gestation during the critical phase of cortical expansion.

The precise mechanisms by which specific HSA21 genes contribute to the neuroanatomical and neurobehavioral phenotypes in DS remain unclear. Aneuploidies cause stoichiometric imbalances in macromolecular complexes, signaling pathways, and gene regulatory networks, and can lead to proteotoxic stress (13). For a subset of genes encoded on an aneuploid chromosome, protein expression levels can be buffered to diploid levels by various post-transcriptional mechanisms (14, 15). For example, in trisomy 21 cells, HSA21 encoded proteins are more likely to be expressed at diploid levels if they participate in macromolecular complexes (16). However, a proportion of genes will not be buffered, resulting in protein levels that increase in proportion to the dosage increase associated with the aneuploidy (15).

The HSA21-encoded protein kinase DYRK1A is essential for normal brain development from flies to humans (17). Loss of function mutations in the *Drosophila* ortholog *minibrain* impair post-embryonic neurogenesis, causing a reduction in brain size (18). In humans, haploinsufficiency of *DYRK1A* underlies the neurodevelopmental disorder MRD7 (OMIM 614104), which is associated with intellectual disability, microcephaly, and autism (19-26). In DS brain tissue, DYRK1A protein levels are expressed at ~1.5-fold higher levels in proportion to HSA21 copy number increase (27). Increased DYRK1A expression has been implicated in disrupting aspects of neurodevelopment and neurodegeneration in DS (28-31). Multiple lines of evidence from mouse models support a causative role for DYRK1A dosage alterations in atypical brain development and intellectual disability (31-36), and point to DYRK1A kinase activity as a potential therapeutic target in DS-associated neuropathologies (28, 37-39).

Despite some success in using mouse models of trisomy 21 to interrogate potential interventions in Down syndrome, anatomical and developmental differences between rodents and humans limit the utility of the mouse in the study of human neurological disorders (40). For example, the rodent central nervous system develops along fundamentally different trajectories involving neural stem cell induction mechanisms distinct from those found in hominids (41, 42). Mouse and rat brains are lissencephalic (smooth), whereas human brains are gyrencephalic (convoluted), with a high surface area thought to originate from proliferative expansion of specific neural progenitor cell subtypes (43, 44) that are poorly represented in rodents (45). The recent failure of a phase II clinical trial to meet primary and secondary endpoints for RG1662, a GABAA inverse agonist (46) that showed promise in a DS mouse model (47, 48), highlights the urgent need for new model systems that more faithfully recapitulate DS neurobiology.

Human stem cell derived cerebral organoids (COs) provide a tractable alternative model system that is amenable to pharmacological and genetic manipulations not possible in humans (49, 50). COs develop into three dimensional structures that mimic important features of the *in vivo* cytoarchitecture found in developing human neocortex (51). Many CO culture protocols begin with the formation of embryoid body aggregates (EBs) that develop regions of radially expanding neuroepithelium upon neural induction, a process that mimics aspects of neurogenesis during human cortical development (52). Many neurodevelopmental disorders arise during these nascent stages of corticogenesis, making COs a useful platform to model complex and uniquely human neurobiological processes and disorders, and to evaluate candidate therapeutics for neuropathologies previously considered untreatable (53).

Large-scale transcriptome and epigenome profiling studies reveal that COs recapitulate many of the key epigenetic and transcriptional programs found in developing human forebrain (54, 55). Post-transcriptional regulatory mechanisms play crucial roles in both neocortex development (56-58) and dosage compensation in DS cells (16). Consequently, trisomy 21 is predicted to have complex effects on proteome remodeling in the developing brain not revealed by steady state mRNA measurements. While comprehensive transcriptome maps of human DS brain tissue in neurotypical and DS individuals have been produced at unprecedented spatio-temporal detail (55, 59, 60), similar efforts to systematically map proteome alterations in DS brain tissue and CO model systems are lacking.

To identify potential mechanisms underlying altered cortical development in DS, we performed ultra-deep, label-free quantitative proteomics at four developmental stages during CO development and differentiation, using human euploid and trisomy 21 induced pluripotent stem cell (iPSC) lines. We also evaluated the proteome-wide effects of the DYRK1A inhibitor, harmine, on the trisomy 21 differentiation trajectory (61). We found that growth in 300 nM harmine led to widespread alterations of the trisomy 21 CO proteome, and partial normalization of functional gene signatures on the protein level for key processes in human cortical development. Integrative analysis of these proteomic changes with a previously described human DS brain mRNA developmental trajectory revealed functional signatures of trisomy 21 dysregulated neuronal differentiation that were partially normalized by harmine treatment.

## EXPERIMENTAL PROCEDURES

### Reagents

DMEM/F12 media, Neurobasal media, Geltrex^®^ low growth factor basement matrix, B27 supplement, N2 supplement, human recombinant insulin, basic fibroblast growth factor (bFGF/FGF2), antibiotic/antimycotic, MEM-NEAA, Corning ultra-low attachment cell culture dishes, and Nunclon Sphera 96-well plates were purchased from Thermo Fisher Scientific (Waltham, MA). TeSR E8 media and Accutase cell dissociation reagent were from StemCell Technologies (Vancouver, BC, Canada). Y-27632 was purchased from Tocris Bioscience (Bristol, UK). Cell culture-grade dimethyl sulfoxide (DMSO), β-mercaptoethanol, Triton X-100 (TX100) and harmine free base were purchased from Sigma-Aldrich (St. Louis, MO). Matrigel^®^ was purchased from Corning (Corning, NY). Immunoglobulin-free BSA and normal goat serum were from Jackson ImmunoResearch. Antibodies were used and sourced as follows: rabbit pAb anti-Ki67 (Abcam #15580). Secondary antibodies were from Jackson ImmunoResearch (West Grove, PA): AlexaFluor488-goat-anti-mouse, Cy3-goat anti-mouse, AlexaFluor647-goat-anti-rabbit; and from Thermo Fischer Scientific: Alexa Fluor 594 goat anti-mouse IgG (H+L) (A-11005) and Alexa Fluor 594 donkey anti-goat IgG (H+L) (A-11058). Click-EdU labeling reagent, 5-ethynyl-2’-deoxyuridine (EdU), was from Molecular Probes. Proteomics: Unless otherwise specified, all reagents used were LC-MS grade and purchased from Thermo Fisher.

### Cell Culture and Maintenance

The iPSC line I50-2 (euploid female), referred to in this paper as D21, was previously described (62), and exhibited a normal karyotype (supplemental Fig. 1A). The iPSC line C2-4 (trisomy of HSA21, female) (63), referred to in this paper as T21, was a generous gift from Gretchen Stein and Chris Link (University of Colorado, Boulder). The C2-4 line exhibited trisomy 21 in karyotype analysis (supplemental Fig. 1B). Genetic stability of the iPSCs was assessed by G-banding karyotype analysis performed at the University of Colorado Molecular Pathology Shared Resource Cytogenetics Core Facility. Metaphase spreads were captured using CCD camera and karyotyping was carried out using the BandView software (Applied Spectral Imaging Inc).

All cell lines were maintained under feeder-free conditions (E8 media with supplements), grown on Geltrex-coated polystyrene 6-well tissue culture plates. For routine passaging, media was aspirated from wells and cells were rinsed briefly with room-temperature, protease-free 0.5 mM EDTA in Dulbecco’s phosphate-buffered saline (DPBS) and then incubated with protease-free 0.5 mM EDTA in DPBS for 5 min at 37 °C. Cells were detached by pipetting media up and down 3-5 times with a 5 mL pipette before being rinsed off the plate and diluted into an equivalent volume of E8 media. Cell suspensions were centrifuged at 400 x g for 2 min, and the cell pellets were resuspended in room temperature E8 media prior to plating. Cells were typically passaged 1:3 when they reached approximately 80% confluence. Cells were treated with 300 nM harmine in 0.003% DMSO, or 0.003% DMSO as vehicle control. To generate the cerebral organoids, we employed the method detailed by Lancaster and Knoblich (52) with some modifications as detailed below. Samples for proteomic analysis were collected as follows. Samples of iPSC cultures were collected after 3 days of culture, prior to EB induction. We initiated EB formation at day in vitro (DIV) 0, and collected EBs at DIV5 prior to neural induction (Fig. 1A). We refer to embryoid bodies that were grown in neural induction media for 5 days as “neurospheres”, which were collected at DIV10. Neurospheres (NS) were embedded in Matrigel at DIV10 for 15 days, followed by 21 days in rotating suspension culture. The COs were collected at DIV45, pooling approximately 10 organoids per replicate.

**Figure 1.**
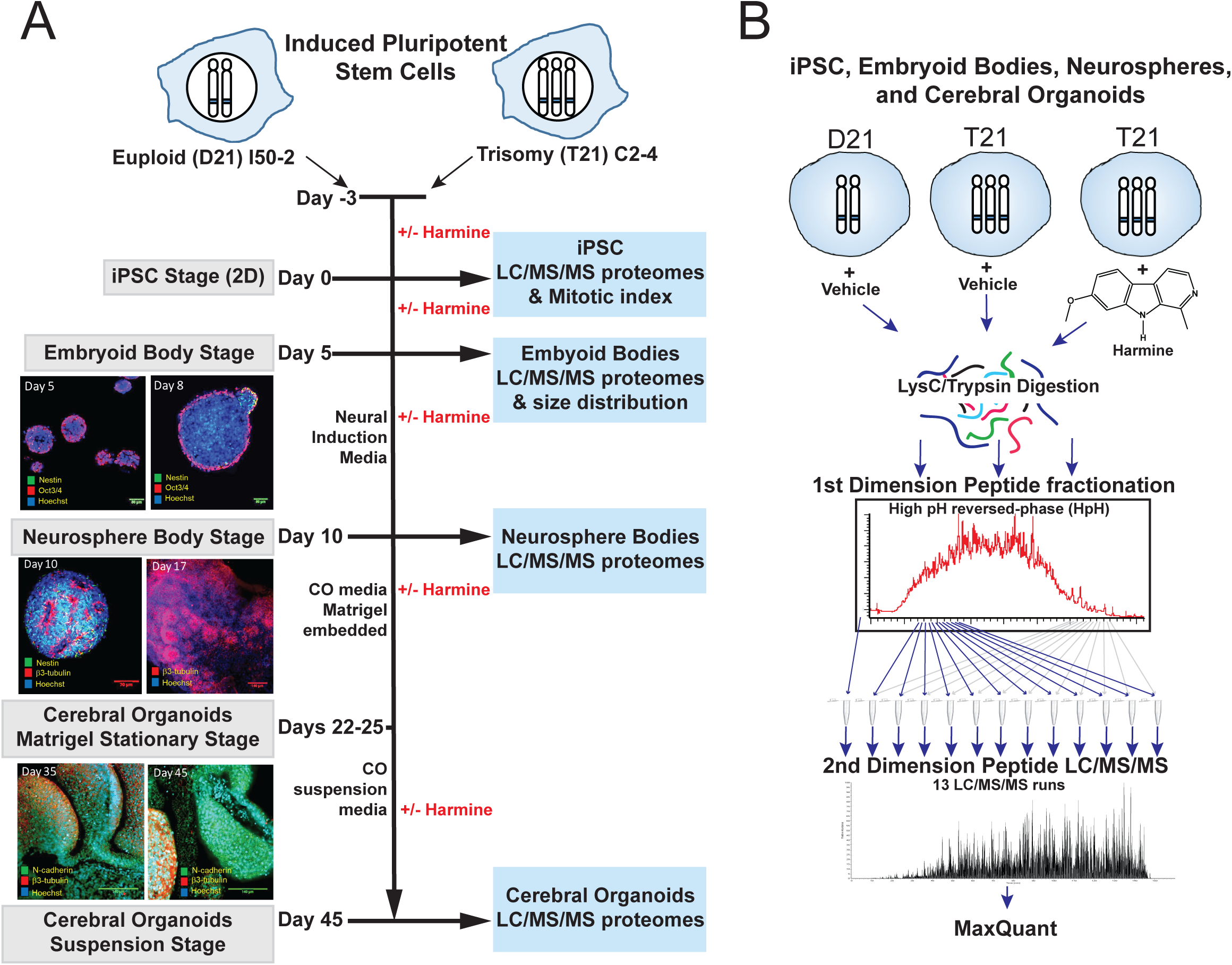
Overview of cerebral organoid generation from human euploid and trisomic 21 iPSCs and proteomic workflow. (**A**) Schematic of cerebral organoid (CO) developmental trajectory, starting from patient derived induced pluripotent stem cells. The trisomy 21 iPSC line C2-4 (T21) was grown and differentiated in the presence of DMSO vehicle (0.003%) or 300 nM harmine (in 0.003% DMSO), and compared to the euploid iPSC line (D21) grown and differentiated in 0.003% DMSO. Four separate stages were compared: iPSC, embryoid body (EB), neurosphere (NS), and CO stages. (**B**) Label-free quantitative proteomics, using multidimensional UPLC peptide fractionation and high-resolution Orbitrap mass spectrometry, was carried out on each of these developmental stages.

Embryoid bodies were seeded as follows, identical to a previously described protocol (64). iPSC colonies were collected as for routine passaging, followed by trituration using a fire-polished Pasteur pipette in complete E8 media supplemented with 50 μM Y-27632 and 4 ng/mL bFGF to obtain a homogenous cell suspension. Cells were counted in a hemacytometer and were seeded at a density of 9,000 cells/well in a volume of at least 100 μl. Cells were fed every other day for 5 days, and monitored visually for successful induction into embryoid bodies and neurectodermal differentiation, as indicated by clearing surface tissue. On the 6th day, embryoid bodies were transferred en masse into neurosphere induction media (DMEM/F12 supplemented with 1x MEM-NEAA, 1x N2 supplement, 1 μg/mL heparin sulfate). The cells were fed every other day for an additional 5-7 days, until neuroectoderm differentiation was apparent. Neurospheres were embedded in Matrigel by putting 10 μl droplets of cold (~4°C) Matrigel onto a 10 cm tissue culture dish on a cold surface. Single neurospheres were collected in 2-3 μl of media and pipetted directly into the cool Matrigel droplet. The plate containing rows of embedded neurospheres was placed into the 37°C cell culture incubator for 10 min to allow the Matrigel to solidify before being covered with 10 mL of primary CO media (1:1 DMEM/F12:Neurobasal, supplemented with 1x B27 [minus Vitamin A – Gibco 12587-010], 1x N2, 1x MEM-NEAA, 3.5 μl/L β-mercaptoethanol, 1x antibiotic/antimycotic). COs were monitored during stationary culture for 10-14 days. COs that exhibited successful neural induction, as indicated by the appearance of multiple neuroectodermal lobes, were transferred to rolling bottle cultures containing mature organoid media (using B27 supplement containing vitamin A - Gibco 17504-044). Media was replaced every 3-5 days until harvest.

### Immunocytochemistry

iPSC growing on Geltrex-coated 1cm diameter round coverslips had media aspirated and were gently washed 1x with 2 mL warm (37°C) DPBS, then fixed with 4% MeOH-free formaldehyde in PBS for 30 min at room temperature. Fixed cells were washed 2x with 1 mL PBS, permeabilized with the addition of 1 mL PBS+0.1% TX100 and gently rocked for 15 min. For cells labeled with EdU (1 μM), fluorescent conjugation was performed by incubating permeablized cells with AlexaFluor-488-azide (10 μM) in the presence of 4 mM CuSO4 and 100 mM ascorbic acid for 30 min at room temperature with gentle agitation. Samples were blocked with 3% IgG-free BSA in PBS+0.1% TX100 for a minimum of 30 min prior to the addition of primary antibodies. All primary antibodies were added at a concentration of 1 μg/mL in fresh 3% BSA/PBS solution and incubated overnight at 4°C. The following morning, primary antibody solution was removed, the samples washed 2x with 1 mL PBS+0.1% TX100, then re-blocked with 3% IgG-free BSA and 1% normal goat serum in PBS+0.1% TX100 for a minimum of 30 min at room temperature. Secondary antibodies were added to the samples at a 1:250 dilution in fresh 3% BSA/1% NGS/PBS solution and incubated for 2 hr at room temp with gentle rocking. Secondary antibody solution was removed and cells were washed twice with 1 mL PBS+0.1% TX100 for 15 min; Hoechst 33342 was added to a final concentration of 10 μg/mL during the first washing step to label cell nuclei. Following the final wash, coverslips were mounted onto glass slides with ~50 μl of Fluoromount G and allowed to dry in the dark overnight before imaging.

### Confocal Microscopy

Images were acquired on a Zeiss 510 laser scanning confocal microscope equipped with an Ar/2 laser for excitation at 458, 477, 488, 514 nm, 2 HeNe lasers for excitation at 543 and 633 nm, 5x/0.15NA, 10x/0.3NA EC Plan Neofluar objectives, a 20x/0.8NA Plan Apo Chromat objective and 40x/1.3NA oil DIC EC Plan Neofluar, 63x/1.4NA oil DIC Plan Apo Chromat 100x/1.4NA oil DIC Plan Apo Chromat objectives, running Zen v.3.0 acquisition software. Image analysis was performed in ImageJ 2.0 (NIH).

### Brightfield Imaging

Images were acquired on an Olympus IX81 inverted widefield microscope through the 4x objective with a Hamamatsu Orca R2 CCD, illuminated by a PriorLumen 2000 light source. For image acquisition of embryoid body formation, serial images of 3 adjacent wells were taken and TIFF images exported for analysis with ImageJ 2.0 (NIH).

### Sample Preparation for Proteomic Analysis

COs were collected and rinsed with DPBS, and rapidly solubilized in 0.5 mL of 4% SDS, 0.1 M Tris (pH 8.5), Lysates were snap frozen in liquid N2 and stored at −80°C for further processing. Lysates were processed for proteomics analysis using a modified version of the filter-aided sample prep (FASP) method (65), described below. Samples were reduced by the addition of 10 mM TCEP, boiled 5 min and incubated at room temperature for 20 min. Reduced sulfhydryls were alkylated with 25 mM iodoacetamide for 30 min in the dark. Approximately 50-100 ug reduced and alkylated total protein was transferred to a washed 30 kD MWCO Amicon Ultra (Millipore) ultrafiltration device and concentrated via centrifugation. SDS was removed by washing three times with 8 M urea, 100 mM Tris pH 8.5, and again three times with 2M urea, 100mM Tris pH 8.5 in a similar fashion. One microgram of endoprotease LysC (Wako, Osaka, Japan) was added and incubated for 3h with rocking at room temperature, followed by one microgram of trypsin (Promega), rocking overnight at room temperature. Tryptic peptides were removed from the ultrafiltration device by centrifugation and stored at −80°C.

### Tryptic peptide fractionation

To reduce peptide sample complexity, 30-50 μg of tryptic peptides were fractionated on a high pH reversed-phase C18 column using a Waters M-class Acquity UPLC. Mobile phases used were (A) 10 mM ammonium formate, pH 10 in LCMS water and (B) 10 mM ammonium formate, pH 10 in 80% (v/v) LCMS acetonitrile. Samples were loaded and washed to baseline on a custom fabricated C18 column (1.8 μm bead size, 120 Å pore size, UChrom C18 (nanolcms) 0.5 mm x 200 mm) at a flow rate of 15 μl/minute equilibrated with 5% B. Peptides were eluted with a gradient from 5% B to 100% over 160 min. A ‘flow-through’ fraction was collected and offline desalted, and fractions were collected for 1 min each, concatenating throughout the gradient for a total of 12 mixed fractions of nearly equal complexity. Fractions were dried by vacuum centrifugation and stored at −80°C.

### Mass spectrometry and data analysis

All fractions for mass spectrometry were suspended in 5% (v/v) acetonitrile, 0.1% (v/v) trifluoroacetic acid in LC/MS-grade water and analyzed by direct injection on a Waters M-class Acquity column (BEH C18 column, 130 Å, 1.7 μm, 0.075 mm x 250 mm) at 0.3 μL/minute using a nLC1000 (Thermo Scientific). Peptides were gradient eluted from 3% ACN, 0.1% formic acid to 20% ACN in 100 min, 20 to 32% ACN in 20 min, and 32 to 85% ACN in 1 min. Mass spectrometry analysis was performed using an Orbitrap Fusion (Thermo Scientific). MS1 were collected at 120,000 FWHM resolution from 380-1500 m/z, 2 × 105 automatic gain control (AGC), 50 ms max fill time, with 20 s dynamic exclusion +/− 10 ppm using the data dependent mode “Top Speed” for 3 s on the most intense ions. MS2 was performed in the linear ion trap with a 1.6 m/z isolation window, 35% HCD collision energy, 1×104 AGC, 35 ms max fill time. All Thermo raw files were processed using MaxQuant (version 1.5.6.5)(66), with MS/MS spectra searched using the Andromeda search engine (67) against a Uniprot human database (reviewed version downloaded April 1, 2015) supplemented with sequences from the CRAPome database (68). The search was limited to peptides with a minimum length of seven amino acid residues, and a maximum of two missed cleavages with trypsin/P specificity, and included variable modifications for methionine oxidation and protein N-terminal acetylation, with carbamidomethyl Cysteine as a fixed modification, with search tolerances of 20 ppm for MS1 precursor ions, and 0.5 Da for MS/MS peaks. Protein and peptide level false discovery rate (FDR) thresholds were set at 1%. The MaxQuant method for label-free quantification (LFQ) was used to generated normalized relative protein abundances, with the match between runs option on, and protein groups quantified using unique and razor peptides. The Perseus software platform (69) (version 1.5.5.3) was used to annotate proteins and filter contaminants from the MaxQuant proteinGroups.txt file. Only protein groups with at least two valid LFQ values (with MS/MS identified MS1 features) over all samples were considered for analysis.

### Data availability

All mass spectrometry raw files, and MaxQuant analysis files (parameters.txt, summary.txt, evidence.txt, proteinGroups.txt and Perseus annotated proteinGroups.txt) are uploaded to the MassIVE repository at UCSD, under the ID “MSV000080549”.

### Experimental Design and Statistical Rationale

Microscopy data from measurements taken in ImageJ were analyzed and graphed in SigmaPlot v13.0 and SPSS v23, using unpaired student’s t-test, ANOVA, reporting 95% confidence intervals. All parametric statistics passed a Shapiro-Wilk test of normality and Brown-Forsyth test of equal variance; values for some measures were log-transformed prior to analysis to ensure normality where appropriate. Data are reported as mean values with error bars depicting the standard error of the mean (SEM). For measurements of phenotypes visualized as individual cells in a larger population, we considered “n” to equal mean values from three random observations of a single coverslip (representing a unique cell culture environment). For observations of intact units made up of multiple cells (embryoid bodies, neurospheres) we considered “n” to be a discrete measurement from each individual multicellular unit. Data points were excluded from analysis where obvious preparation failures were evident (i.e. lack of specific staining, cells imaged were not fixed in mounting media). For proteomic profiling, biological replicates were defined as separate plates at the iPSC stage, and independent pools of multiple spheroids at the EB, NS and CO stages.

### Methods for Figure 2

Principal components analysis (PCA) of the relative protein abundance values (MaxQuant LFQ values) was performed using the R package FactoMineR (70). The proteins most highly correlated and anticorrelated with PC1 and PC2 were selected using the dimdesc function (p < 0.01 and absolute value of Pearson correlation coefficient > 0.7). For enrichment analysis, the UniProt identifiers were converted to gene symbols, removing duplicate identifiers. The Fisher exact test was used to test for significant enrichment of these sets against four libraries downloaded from the EnrichR and ARCHS4 projects (71-73): ARCHS4_Tissues, GO_Molecular_Function_2017, Jensen_COMPARTMENTS, and Reactome_2016. We used custom-built R code to perform the enrichment analysis to enable control of the set of genes used as the enrichment background, which avoids bias introduced by whole genome background gene sets used by the online EnrichR tool. The background used for performing the Fisher exact test was the complete set of gene symbols that mapped to proteins in the PCA analysis. The heatmap of proteins correlated with PC1 and PC2 in Fig. 2B was generated using the morpheus R interface package (MIT Broad Institute).

**Figure 2.**
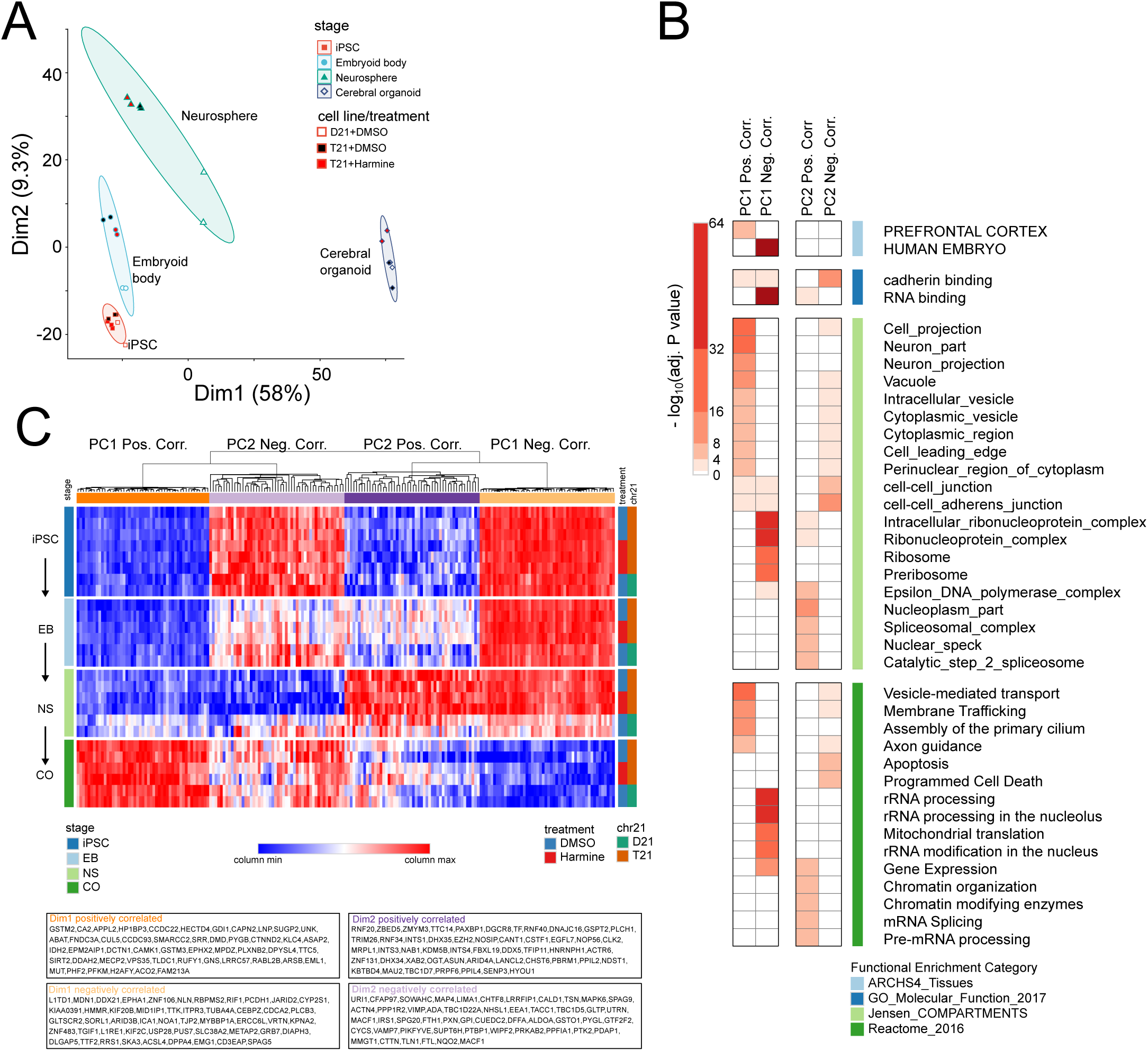
Proteome changes in the developmental trajectory reflect a shift from embryonic to prefrontal cortex specific expression. (**A**) Principal component analysis of label-free intensities (LFQ) shows that the transition to organoids, rather than trisomy 21 or harmine treatment, contributes the most to the variance in protein abundance. The ellipses indicate the 80% confidence regions. (**B**) Functional enrichment analysis of those proteins most highly correlated or anti-correlated with the first and second principle components (PC1 and PC2). Proteins were selected based on a Pearson correlation of > 0.7 and p-value < 0.01. Fisher exact test enrichment was performed against the following gene set libraries: ARCHS4 Tissue, Gene Ontology (GO) molecular function, Reactome and subcellular localization annotations. (**C**) Expression heatmap of the top 50 proteins most correlated and anti-correlated with the first two principal components. Samples are shown in rows and column normalized protein LFQ in columns, with the developmental stage indicated by the left side bar and the HSA21 status and treatment condition shown in the right sidebars.

### Methods for Figure 3, Supplemental Figure 4 and Table I

Protein LFQ values were considered for differential analysis if the total number of missing intensities for that protein group in each stage was less than three. Differential analysis of the LFQ protein abundances for the two comparisons, T21 vs. D21 and T21+DMSO vs. T21+harmine, was performed by using the empirical Bayes modeling framework in the R package, limma (74, 75), with the resulting p-values corrected for multiple-hypothesis testing using the Benjamini-Hochberg method (76). Gene set enrichment analysis was performed for the proteins showing differential expression in the T21 vs. D21 and T21 +/− harmine comparisons (FDR < 0.1). Differential analysis statistics and annotations for all quantified proteins in the trajectory are reported in supplemental Table 2, and CO stage differential analysis for the set of markers shown in Fig. 4B are reported in supplemental Table 3.

**Figure 3.**
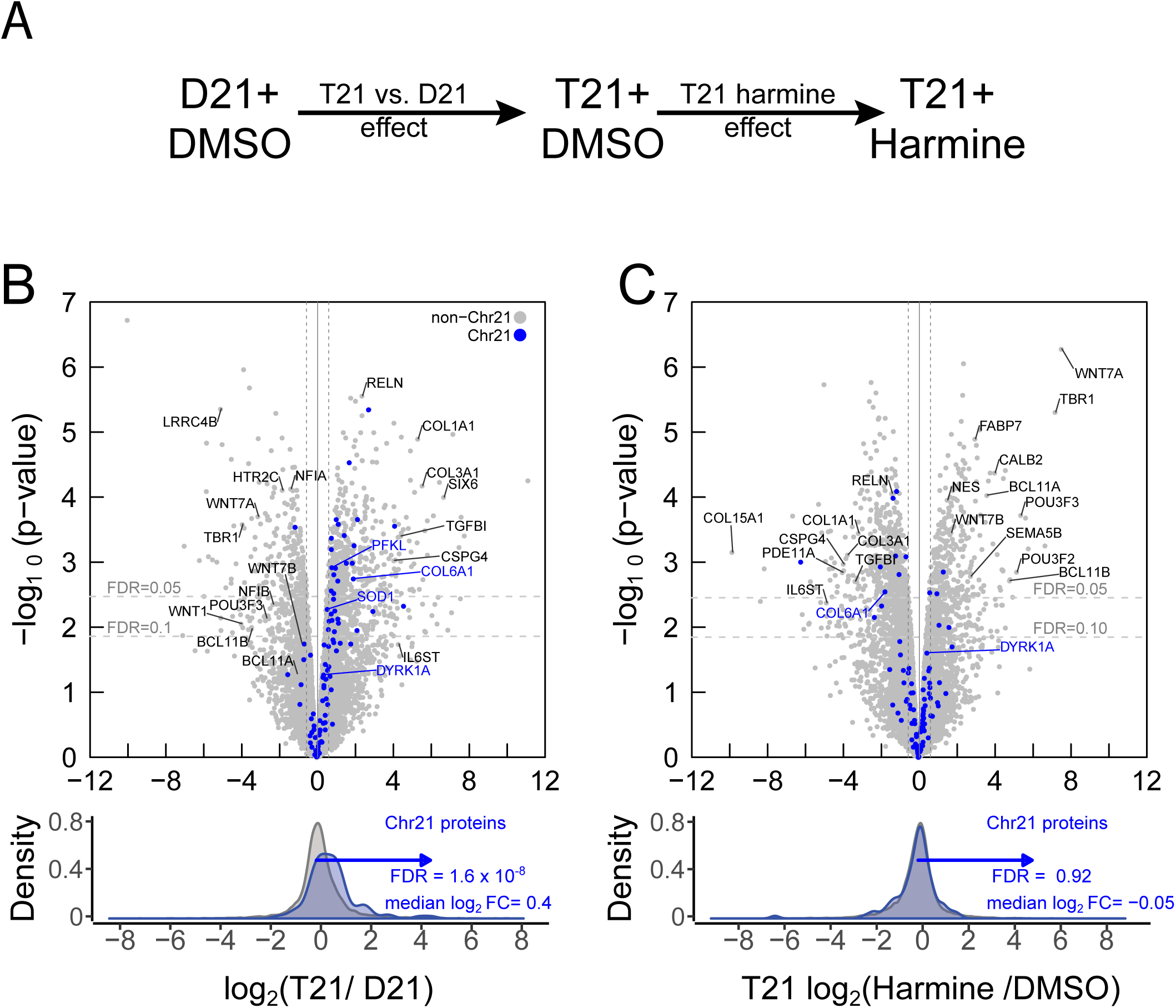
Differential protein expression analysis for T21 vs D21 and harmine-treated T21 cerebral organoids. (**A**) Diagram of comparisons that were tested in the differential expression analysis in each of the four stages, and shown for the CO stage in panels (**B**) and (**C**). The arrows represent the estimated log2 ratios (coefficients) for the two comparisons. For each coefficient, the arrow points toward the numerator in each log2 ratio. (**B**) Volcano plot of the log_2_ ratio of label free abundances (LFQ) vs. −log_10_(p-value) from an empirical Bayes moderated t-test of the CO T21+DMSO (C2 line) vs. D21+DMSO (I50-2 line) comparison, with HSA21 encoded proteins indicated in blue. The horizontal dotted lines indicate p-value thresholds corresponding to the indicated false-discovery rates as determined by multiple testing correction with the Benjamini-Hochberg corrected p-values. The vertical dotted lines correspond to 50% change in abundance from a ratio of one. Positive log_2_ ratios indicate higher expression in the T21 COs relative to D21 COs. (**C**) Volcano plot of the CO T21+300 nM harmine vs. T21+DMSO treatment comparison. Positive log_2_ ratios indicate higher expression in the T21+harmine condition.

**Figure 4.**
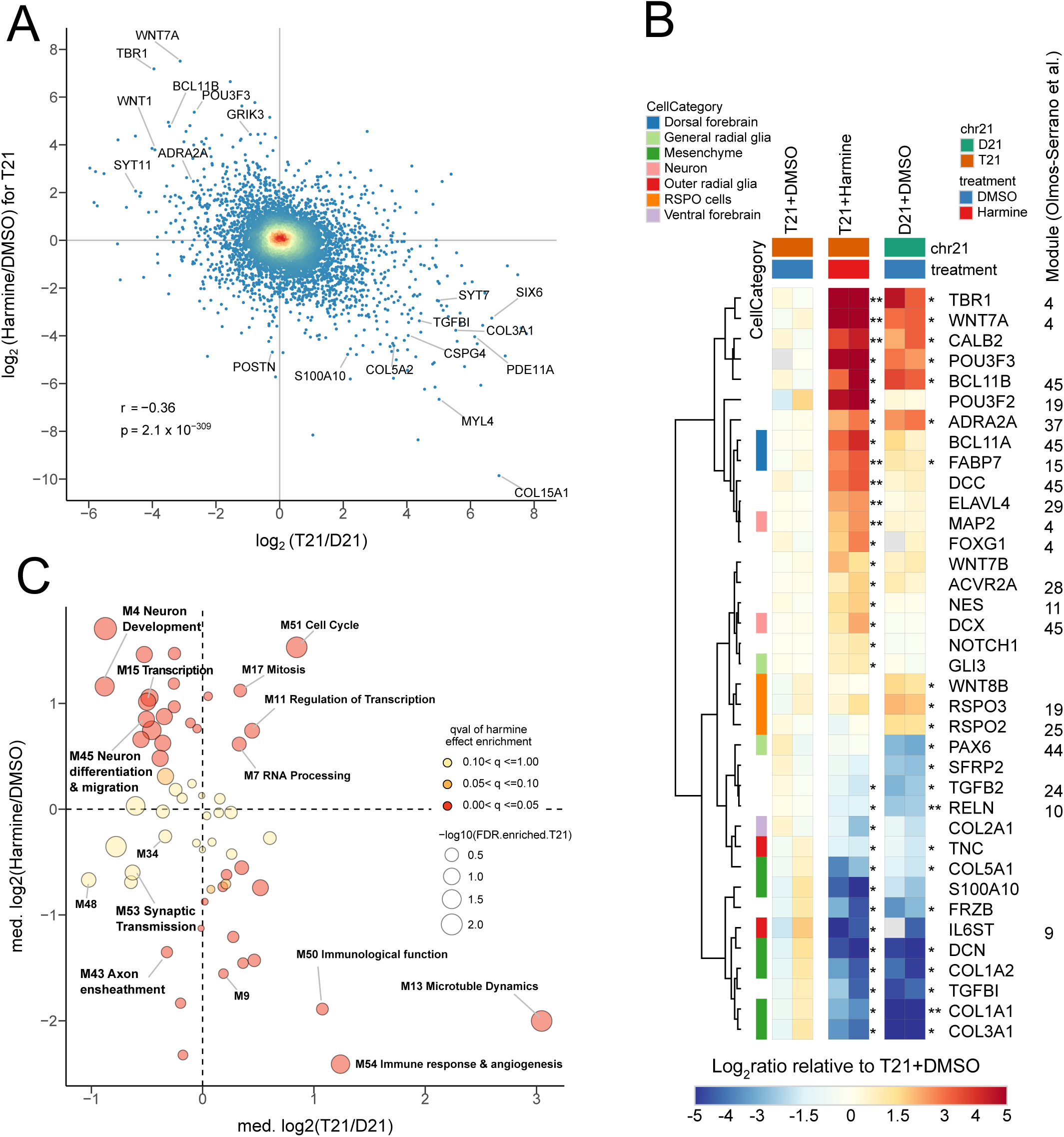
Harmine treatment normalizes trisomy 21 associated expression differences, which overlap with gene expression modules identified as dysregulated in trisomy 21 developing human brain. (**A**) Scatter plot of the log2 abundance ratios from the T21/D21 and T21 +/− harmine comparisons, with color indicating the point density. Shown is the Pearson correlation coefficient (r) and the p-value from significance testing of the correlation. (**B**) Protein expression (LFQ) heatmap for select proteins involved in forebrain development for T21, T21+harmine and D21 COs. The abundances for each protein are normalized to the average abundance in the T21 CO samples. Proteins used as markers or uniquely associated with particular cell types in the developing brain are annotated with their respective cell types in the left sidebar. Hierarchical clustering of relative expression patterns was used to order the rows. Asterisks annotating the second and third column groups indicate statistical significance of the respective comparisons in the empirical Bayes analysis shown in Fig. 3C and 3B, respectively. Proteins that map to functional modules identified in Olmos-Serrano et al. (2016) are annotated with the module number on the right. (**C**) Functional enrichment of differentially expressed proteins in COs was assessed independently for both the T21/D21 effect (X-axis) and the harmine effect (Y-axis), using the functional gene set modules identified by Olmos-Serrano et al. (2016) in post-mortem human brain tissue from Down syndrome individuals and age-matched euploid controls. The Wilcoxon rank sum test was used for threshold-free significance testing of the respective t-statistic for each effect. Modules showing enrichment for differentially expressed proteins (FDR < 0.1) are plotted with the median t-statistic for both T21 vs. D21 and harmine effects. Functional annotations are from Olmos-Serrano et al. (2016) for indicated modules. Color indicates the Benjamini-Hochberg corrected p-value for enrichment of the harmine effect, and size indicates the corresponding corrected p-value for the T21 vs. D21 effect.

**Table I:** Number of Differentially expressed proteins at q < 0.1 at each differentiation stage^a^.

| stage | T21+DMSO vs. D21+DMSO<sup>b</sup> |  | T21+Harmine vs. T21+DMSO<sup>c</sup> |  |
| --- | --- | --- | --- | --- |
|  | T21 high | T21 low | T21 Harmine Up | T21 Harmine Down |
| iPSC | **752** (IPSC_T21HIGH) | **390** (IPSC_T21LOW) | **0** | **1** |
| EB | **1028** (EB_T21HIGH) | **993** (EB_T21LOW) | **1** | **0** |
| NS | **1101** (NS_T21HIGH) | **761** (NS_T21LOW) | **0** | **0** |
| CO | **946** (CO_T21HIGH) | **498** (CO_T21LOW) | **615** (CO_T21_HarmineUP) | **883** (CO_T21_HarmineDN ) |

a: q-value refers to B.H. corrected p-values to control false discovery rate at 10%.
b: total proteins passing threshold (high or low in T21+DMSO vs D21+DMSO)
c: total proteins passing threshold (high or low in T21+Harmine vs T21+DMSO)

### Methods for Figure 4

Functional enrichment using the Wilcoxon rank sum test was assessed independently for both the T21 vs. D21 effect (x-axis) and the harmine effect (y-axis), using the functional gene set modules identified by Olmos-Serrano et al. 2016 in post-mortem human brain tissue from Down syndrome individuals and age-matched euploid controls (60). The moderated t-statistics from the empirical Bayes analysis in Fig. 3 were used in the Wilcoxon rank sum test. Modules showing enrichment for differentially expressed proteins are plotted with the median log2 abundance ratio for both T21 vs. D21 and T21+harmine effects. Functional annotations and labels for modules in Figure 4C were defined in Olmos-Serrano et al. 2016 as follows: M4 Neuron Development, M7 RNA Processing, M9 Homeostatic Mechanisms, M11 Regulation of Transcription, M15 Transcription, M17 Mitosis, M34 Synaptic transmission/regulation, M43 Axon ensheathment, M45 Neuron differentiation/migration, M51 Cell Cycle, M53 Synaptic Transmission, M54 Immune response & angiogenesis. The complete set of enrichments are listed in supplemental Table 4.

### Methods for Supplemental Figure 5,6

Enrichment analysis was performed with the Fisher exact test as in Fig. 2B, using cell type specific gene sets assembled using supplemental tables from previous studies (54, 77), and the indicated libraries downloaded from the EnrichR and ARCHS4 projects (71, 72, 78). Complete enrichment results are listed in supplemental Tables 5 and 6.

### Methods for Figure 5

To generate the network, we used Cytoscape (v3.5.1), and performed a STRING protein query through the “Import Network from Public Databases” dialog, entering all differentially expressed proteins in harmine treated T21 COs (q < 0.1), using confidence score cutoff equal to 0.55 and maximum additional interactors equal to zero. The log2 ratios for the T21 vs. D21 and T21+harmine effects were then mapped to node border color and node fill color, respectively. This network was then clustered using MCODE (79) within the clusterMaker app (80). MCODE parameters were as follows: include loops, unchecked; Degree cutoff, 2; Haircut, checked, Fluff, checked; Node score cutoff, 0.2; k-Core, 2; Max Depth, 100; Create new clustered network, checked. The resulting networks were then exported as an svg file. Functional enrichment analysis was used to annotate each cluster with the most significant gene ontology function. The edge thickness and grayscale indicates the confidence in interaction (81).

**Figure 5.**
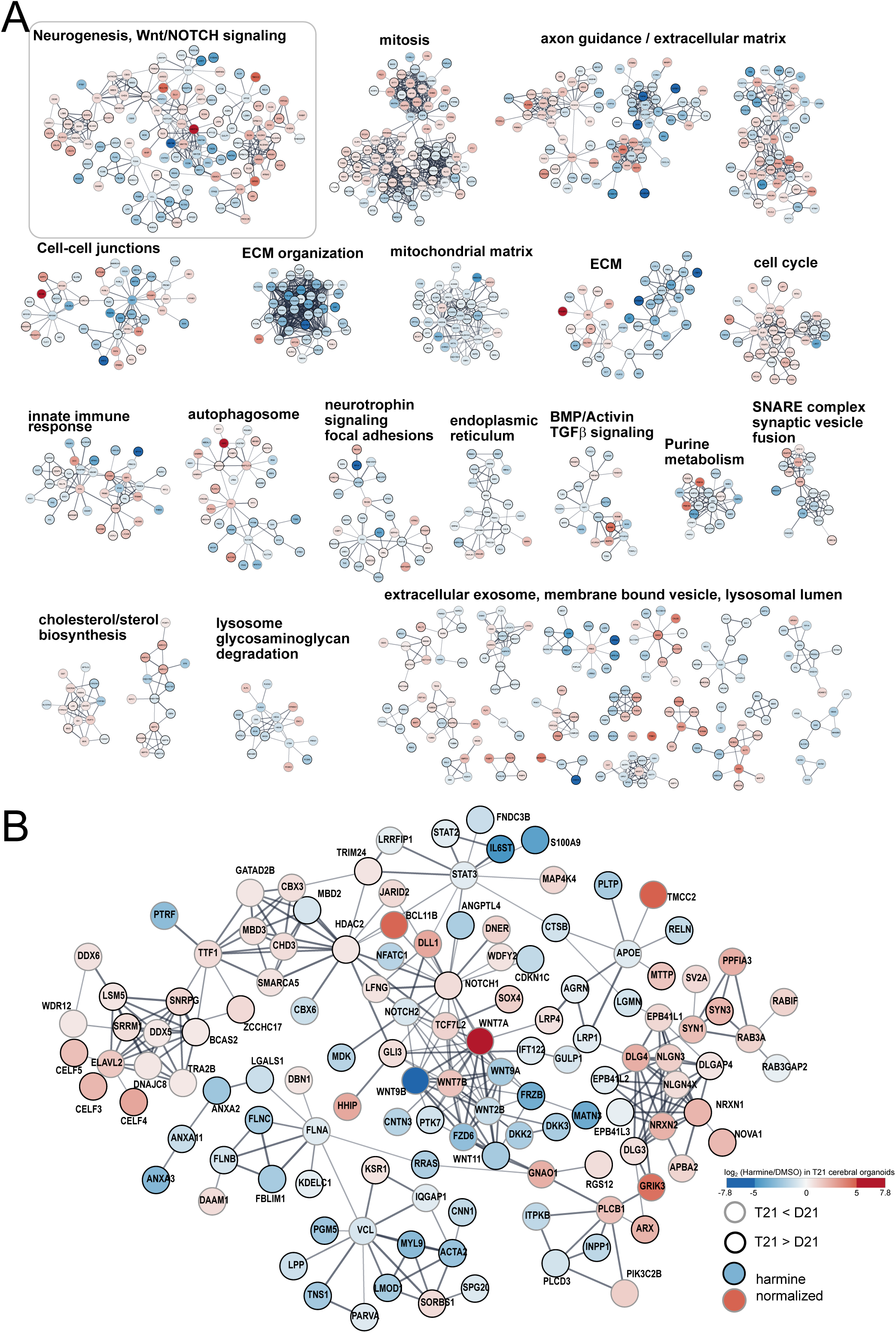
Protein-protein and functional interaction networks of harmine-modulated proteins in COs. (**A**) Differentially expressed proteins in harmine treated T21 COs (q < 0.1) were mapped onto STRING protein interaction networks using the STRING Cytoscape app. The resulting network was clustered using MCODE to find densely interconnected subnetworks, which were then functionally enriched to annotate each cluster with the most significant gene ontology function. The edge thickness and grayscale indicates the interaction confidence level. **(B)** The subnetwork from panel A for a subnetwork associated with WNT signaling and neurogenesis. The node color is mapped to the harmine effect size, log2(T21 harmine / T21 DMSO), and the node border color indicates the direction of change with respect to the T21 vs. D21 effect, log2(T21 DMSO / D21 DMSO).

## RESULTS

### Harmine effects on mitotic index and embryoid body size for euploid and trisomy 21 lines

We used two human iPSC lines grown in feeder-free culture for proteomic and phenotypic analyses: one line was derived from a female individual with Down syndrome (C2-4 line, referred to hereafter as T21) (63) and a second was derived from an unrelated female euploid individual (I50-2 line, referred to hereafter as D21) (Experimental Procedures). To modulate DYRK1A activity, we used the β-carboline alkaloid, harmine, based on reports that DYRK1A inhibition and knockdown was reported to rescue impaired neurogenesis in DS iPSC-derived neural progenitor cell cultures (82). Harmine has also been shown to increase proliferation in euploid neural progenitor cells (83), and remains one of the most selective DYRK1A inhibitors to date (84).

To identify an optimal concentration of harmine for proteomic analysis of the T21 organoid differentiation trajectory, we examined EB size after continuous exposure to a range of harmine concentrations, from the iPSC to the EB stage. In the absence of drug, EBs produced from T21 iPSCs were ~61% (s.e.m. = 17%) smaller relative to D21 EBs (supplemental Fig. 2A). Below 1 μM, harmine produced a dose-dependent increase in T21 EB size, with no significant change for D21 EBs. At concentrations above 1 μM, harmine led to reduced EB size in both T21 and D21 lines, indicating a negative effect on viability and/or growth rate. Based on these observations, we tested the effect of 300 nM harmine on proliferation in the iPSC lines. The T21 iPSC line exhibited a lower mitotic index relative to D21; treatment with 300 nM harmine resulted in an increased mitotic index to the level found in untreated D21 cells (supplemental Fig. 2B), consistent with previous studies showing normalization of DS related mitotic phenotypes by DYRK1A inhibition (82). In HeLa cells, equivalent harmine concentrations (330 nM) have been reported to inhibit phosphorylation of the DYRK1A substrate, SF3B1, on Thr434 by 30-50% (85). Submicromolar harmine has been shown to inhibit DYRK1A-dependent phosphorylation events in cells (61), and to attenuate axonal growth in mouse primary neurons overexpressing Dyrk1a (86). Based on these results, we used 300 nM harmine, present throughout all differentiation stages, from iPSC to CO, for the proteomic analysis of the CO developmental trajectory (Fig. 1A).

### Proteomic analysis of developmental trajectories for euploid, trisomy 21 and harmine treatment

To identify proteins that change in abundance during the differentiation process, and show differential expression with harmine treatment, we generated quantitative proteomes at multiple stages along the CO developmental trajectory (Fig. 1A) (Experimental Procedures). All lines were grown in parallel to match culture conditions (T21 + DMSO, T21 + 300 nM harmine, and D21 + DMSO), using a modified version of the Lancaster protocol (52) similar to that used in a recent Zika virus study in COs (64) (Experimental Procedures). We collected three biological replicates at the iPSC stage and two biological replicates at the spheroid stages (EB, neurosphere, CO), pooling multiple spheroids in each of the last three stages to obtain sufficient material for high-depth label-free proteomic profiling. Biological replicates in this experiment refer to independent cultures grown in parallel. We found that 300 nM harmine treatment led to the systematic failure of euploid (D21) iPSC-derived CO formation; consequently, D21 harmine-treated samples were not collected for any stage in this experiment.

We used an optimized offline peptide fractionation method to simplify peptide mixtures for LC/MS/MS proteomic analysis on a quadrupole/linear ion trap Orbitrap mass spectrometer (Fig. 1B, Experimental Procedures) (87). Extraction of label-free quantitation (LFQ) values, which are normalized relative protein abundance measures, was performed with MaxQuant software (66). In total and across all stages and samples, after applying a protein and peptide FDR of 1%, we identified 12,869 proteins (identified by at least two MS/MS) and quantified 11,225 proteins (proteins quantified in fewer than two samples across all 26 samples were not counted). Approximately ~80% of the quantified proteins were common to all four stages (supplemental Fig. 3A). The organoid stage showed the largest number (399) of proteins uniquely expressed in a single stage. Correlation between biological replicates was very high, with Pearson correlation coefficients 0.96-0.99 (supplemental Fig. 3B).

To examine the dominant trends in protein abundance dynamics over the four developmental stages, we performed principal component analysis (PCA) on the protein LFQ values (Experimental Procedures). Samples clustered by stage in the PCA scores plot (Fig. 2A), indicating that variability between the two cell lines is secondary to the larger variation associated with the developmental process. Most of the total dataset variance is captured in the first principal component, PC1 (58%), which is dominated by the developmental transition from NS to CO, during which neurogenic precursors multiply and differentiate into neurons (54). The second principal component (PC2) captures the developmental transition from iPSC to NS, and accounts for a relatively minor proportion (9.3%) of total dataset variation. These results are consistent with previous reports showing that neurogenesis and neuronal differentiation in COs and iPSC derived neurons involve highly dynamic remodeling of both the transcriptome and proteome (55, 88).

### The developmental trajectory is characterized by temporal patterns of protein expression related to loss of pluripotency and acquisition of neuronal cell fates

To gain further insight into the biological processes associated with these transitions, we performed functional enrichment analysis on the proteins most highly correlated and anti-correlated with the first two principal components, PC1 and PC2 (Fig. 2B, C). We tested for non-random overlap of these proteins with gene set libraries from the EnrichR project (71-73), which includes libraries related to molecular function (89-91), subcellular compartments (92), and tissue-specific gene sets from the ARCHS4 libraries from EnrichR. The ARCHS4 libraries contain gene sets derived from large-scale co-expression analysis that identifies highly co-varying sets of mRNAs over thousands of different perturbations and genetic backgrounds in human cell lines and tissues. Enrichment analysis against these libraries can be used as an unbiased method to infer gene function, protein-protein interactions, and human tissue expression patterns (73) (Experimental Procedures).

Proteins positively correlated with PC1 were characterized by low expression in the iPSCs, EBs and NSs, and high expression in COs (Fig. 2C). In this group, we found enrichment for terms related to human brain specific expression (Fig. 2B, column “PC1 Pos. Corr.”, ARCHS4_Tissue library), with prefrontal cortex being the most significant (top row, left column Fig. 2B, adj. p-value 8.8 × 10^−6^, and supplemental Table 1). We also observed significant enrichment in proteins found in postsynaptic density structures (not shown, adj. p-value = 2.3 × 10^−4^), and other neuronal subcellular structures and functions (Fig. 2B). Proteins enriched in this category included DCX, which is important for neuronal migration and cortical lamination, and NCAM1, which is involved in neurite extension (93). Significant enrichments for subcellular structures important during neurogenesis were consistently found. These included cell-cell junctions, which are critical for the regulation of neuroepithelial cell polarity during early forebrain development (94). Intriguingly, highly significant enrichment was found for functions related to primary cilium assembly. The term “*Assembly of the primary cilium”* from Reactome showed a high degree of overlap (56 out of 187) with PC1 correlated proteins (adj. p-value = 2 × 10-11). This category included 15 intraflagellar transport complex proteins, four BBSome subunits (BBS2, BBS7, BBS9 and LZTFL1), and PAFAH1B1. These proteins are involved in primary cilia assembly and trafficking, and have essential roles in morphogen signaling and patterning during brain development. Disruption of many of these genes cause severe neurodevelopmental defects in humans (95). Primary cilia play an important role in the proliferation and differentiation of neuroepithelial cells and apical radial glial cells, which undergo dramatic proliferative expansion during brain development, and are present in COs (96). Both of these cell types have a primary cilium anchored in the apical plasma membrane that projects into the ventricular lumen. Primary cilia are involved in the establishment and maintenance of apical-basal polarity of radial glial cells in the developing brain (97). Expression in this group of proteins sharply increased in COs, suggesting that cell types with primary cilia increase in number during organoid maturation.

The PC1 anti-correlated proteins, which show high expression in the first three stages and low expression in the CO stage, are enriched in functions related to RNA binding, rRNA processing and mitochondrial translation. Consistent with loss of pluripotency and commitment toward neuronal identities over the course of the developmental trajectory, we found highly significant enrichment in markers of human embryonic tissue (Fig. 2B) and neuronal progenitor cells (adj. p-value = 2.3 × 10-10, ARCHS4_Tissue term “*NEURONAL PROGENITOR CELLS*”, supplemental Table 1). Many of the PC1 anti-correlated proteins that map to this progenitor cell type term are involved in regulating the cell cycle and mitotic spindle (e.g. PLK1, CENPF, CCNB2 and AURKA), and include ASPM, which is associated with microcephaly in humans when both copies are disrupted (98). We found two other microcephaly-associated genes that were associated with the set of PC1 anti-correlated proteins, WDR62 and CENPJ, although these were not found to be enriched in these same functional categories. Together, ASPM, WDR62, and CENPJ are involved in spindle orientation in apical progenitors (99), and regulate centriole duplication and the timing of apical progenitor delamination (100).

Proteins correlating with PC2 show differential expression patterns specific to the neurosphere stage. The expression for PC2 positively correlated proteins (PC2 Pos. Corr. in Fig. 2B) increased transiently at the neurosphere stage. Significantly enriched functions for this group are related to the organization of chromatin and heterochromatin domains (e.g. EZH2, RNF20, CBX5, ATRX, EHMT2), and spliceosome function (e.g. U2AF1, SRSF1, PRPF6, DDX5). Proteins anti-correlated with PC2, which decreased transiently at the neurosphere stage, are enriched in functions associated with cell-cell adherens junctions and cadherin binding, including CDC42, CDC42BPA, CSNK1D, ITGB1 and SHROOM3. Adherens junctions play a crucial role in polarity and orientation of apical progenitors (apical radial glia) in the developing forebrain(101). In the developing mouse telencephalon, depletion of CDC42, which regulates the asymmetric localization of the Par complex in apical progenitors, leads to the loss of apical-basal polarity and a shift from apical to basal progenitor fates (102).

To summarize, we found that the greatest variation in protein abundance is associated with the last two developmental transitions (NS and CO). During these stages, neurogenic precursor cells undergo proliferative expansion, and early born neurons begin differentiation and maturation, leading to a diversification of multiple cell types within the organoid. Functional analysis of changes in proteins at the neurosphere stage revealed alterations in chromatin structure, cell polarity, and cell-cell interactions, suggesting important roles for these processes during neural induction. At the CO stage, we found coordinated upregulation of proteins with functions related to neuronal differentiation and primary cilia assembly, and downregulation of proteins involved in maintaining pluripotency.

### Stage-specific differential expression analysis

Having established the primary sources of proteome variation and associated functional signatures, we next performed a differential expression analysis to identify proteins that showed differences between the T21 and D21 lines and responded to harmine treatment at each differentiation stage. We used an empirical Bayes moderated t-test method that provides greater statistical power to detect differentially expressed genes than the unmoderated Student’s t-test, particularly when sample numbers are low (75). This method has been validated for proteomics data (103), and has been successfully used in large-scale differential proteomics experiments (104-106). Differential analysis was performed using custom R scripts that employ the LIMMA modeling framework (74) (Experimental Procedures).

At each stage, we detected 500-1000 differentially expressed proteins in the T21+DMSO vs. D21+DMSO comparison, hereafter referred to as the T21 vs. D21 effect (Table I, Fig. 3A). We found that proteins encoded on HSA21 were expressed, on average, at 30-50% higher levels in T21 across the four stages, relative to the euploid control, consistent with the increased gene dose associated with the extra copy of chromosome 21 (Fig. 3B and supplemental Fig. 4A, C, E). However, a number of proteins deviated from the expected 1.5-fold increase, consistent with recent proteomic studies in trisomy 21 and other aneuploid human cell lines (15, 16).

Interestingly, at the iPSC, EB and NS stages, we detected only two differentially expressed proteins with harmine treatment, that is, in the T21+DMSO vs. T21+harmine comparisons, hereafter referred to as the T21+harmine effect (Table I). In contrast, harmine treatment led to large-scale changes in protein abundance in T21 COs, resulting in 1498 differentially expressed proteins for the CO T21+harmine effect (q < 0.1) (Fig. 3C, Table I). Examination of the log2 ratio distributions for the T21+harmine effect, at each stage in the T21 line, revealed that the iPSC, EB and NS log2 ratios exhibit much less dispersion around zero, compared to the CO T21+harmine effect (supplemental Fig. 4B, D, F, G). This suggests that harmine treatment resulted in fewer large-magnitude effects on protein expression at earlier stages, where cell type heterogeneity is lower compared to the CO stage.

We next examined the proteins with differential expression specific to the CO stage, where harmine treatment exhibited the most dramatic effect on abundance (Fig. 3C). We found that a number of the proteins with the lowest expression in T21 COs, relative to D21 COs, were Wnt ligands (e.g. WNT7A, WNT7B, WNT1) and transcription factors involved in the specification of laminar identity in post-mitotic neurons (e.g. TBR1, BCL11B)(107-109). Strikingly, harmine treatment increased the expression for these proteins in T21 COs to near D21 levels (Fig. 3C). Conversely, many extracellular matrix proteins were expressed at higher levels in T21 COs, relative to D21 COs. For example, Reelin (RELN), an extracellular glycoprotein involved in regulating neuronal migration during cortical lamination, was ~5-fold higher in T21 COs, and decreased by 2.6-fold with harmine treatment. Other ECM proteins that exhibited this trend (higher T21 vs. D21 CO expression and reduction by harmine) included *COL1A2, DCN, and COL5A1,* proteins recently found to be uniquely expressed in a mesenchymal cell type identified in human fetal brain and COs by single cell RNA-seq (54). In this same study, COL5A1 was found in the periphery of CO cortical regions, colocalized with ECM embedded collagen fibers, suggesting that T21 COs contain an increased population of mesenchymal cell types and/or upregulated ECM secretion that is normalized by harmine treatment. Dysregulated extracellular matrix remodelling would be expected to disrupt neuronal migration and self-renewal of neurogenic precursor cells in the developing brain (110).

Overall, we found a large number of proteins with differential expression in the T21 vs. D21 comparison that changed in abundance toward D21 expression levels upon harmine treatment. Of the 1,444 proteins that were differentially expressed in the T21 vs. D21 CO comparison (Table I), an unexpectedly large number, 425 (29%), changed significantly with harmine (p < 2.2 × 10-16, odds ratio = 3.1, Fisher exact test). Of the 946 proteins that were high in the T21 COs relative to D21 COs, 3.3-fold more proteins decreased with harmine than increased (226 vs. 68) Conversely, of the 498 proteins that were low in the T21 COs, 1.4-fold more proteins increased than decreased (77 vs. 54) (supplemental Fig. 5A-C). To test whether this “harmine-normalization effect” was evident on a global level, we performed a correlation analysis between the two differential expression effects, using the LFQ values in the T21+DMSO COs as a baseline for comparison (Fig. 4A). The T21+harmine and T21 vs. D21 effects were highly correlated, with a Pearson correlation coefficient of −0.36 (p-value = 2 × 10^−309^), suggesting that harmine treatment normalizes a significant fraction of the proteome disrupted by trisomy 21. Expression levels in the CO stage, for a selected subset of proteins involved in neocortex development, are shown (Fig. 4B), relative to the T21+DMSO average LFQ values to facilitate visualization of normalization toward D21 levels (data reported in supplemental Table 3).

### Concordance between human DS brain signatures and dysregulated CO protein expression

To clarify the potential functional ramifications of differential expression in the CO comparisons, we mapped differentially expressed proteins onto a recently described transcriptome analysis of developmental stages for normal and Down syndrome human brains, spanning fetal to adult stages (60). In that study, the WGCNA algorithm (111) was used to identify gene modules with highly correlated temporal mRNA expression patterns, many of which were discordant between euploid and Down syndrome brains. Our analysis assumes that discordant mRNA expression will be reflected on the protein level, and should therefore capture dosage-sensitive proteomic changes related to trisomy 21 that are not under post-transcriptional control. In this way, we can determine whether the impacts of trisomy 21 observed in the proteome are consistent with dysregulated signatures observed in human DS brain.

We tested whether the proteins that map to a given DS brain module have significantly increased or decreased expression in each of the two CO comparisons used in this study, T21 vs. D21 and T21+harmine, adapting a previously described statistical method for threshold-free annotation enrichment (112). The method uses a two-sided Wilcoxon rank sum test on the empirical Bayes t-statistics from the differential analysis used in Figure 3, producing a position score and an adjusted p-value for each DS brain module that indicates whether there is significantly increased or decreased protein expression within the module (Experimental Procedures). This analysis enabled us (i) to identify proteins in COs with functional overlap with the dysregulated gene modules in human DS brain, and (ii) to visualize module-specific biases in the two treatment effects (Fig. 4C), analogous to the two axes of the scatter plot in Figure 4A.

We found many DS brain modules with significant shifts in median protein log2 ratios (for both CO effects) that mirrored the correlation structure of the protein log2 ratio scatter plot in Figure 4A (Fig. 4C). That is, many co-expressed gene modules found in DS brain tissue were associated with significant expression differences in the T21 vs. D21 and T21+harmine comparisons. For example, the M4 and M45 modules, enriched in genes involved in neuron differentiation and development, showed significant expression shifts for the T21 vs. D21 and the T21+harmine comparisons, with an overall shift toward lower T21 expression, relative to D21, and increased T21 expression upon harmine treatment (Fig. 4C, supplemental Table 4). A complementary enrichment method, using the differentially expressed protein sets (q < 0.1) across all stages, confirmed that the CO stage showed the most highly significant enrichment in DS brain modules (supplemental Fig. 6A).

The M45 module mapped to 36 differentially expressed proteins that were up-regulated by harmine in the T21 COs, including BCL11A, BCL11B, CHD4 and CHD3 (Fig. 4B, C, supplemental Fig. 6A, supplemental Table 4). BCL11A and BCL11B are paralogous subunits of the BAF complex that are critical for neuron subtype specification (113). Loss of function mutations in BCL11A cause intellectual disability, which has been attributed to transcriptional dysregulation during cortical development (114). CHD4 and CHD3 are mutually exclusive subunits of the chromatin remodeling NuRD complex, which regulates neural progenitor proliferation and layer specification of differentiated neurons during cortical development (115).

The M4 module mapped to 33 differentially expressed proteins that were upregulated by harmine in the T21 COs; these included WNT7A, TBR1 and FOXG1 (Fig. 4B). WNT7A was one of the largest magnitude changes induced by harmine treatment, and plays a crucial role in forebrain development and synaptic organization (116). *WNT7A* mRNA expression was previously reported with low expression in DS iPSC-derived neural progenitors, relative to an isogenic euploid control (82). In the same study, knockdown of *DYRK1A* expression rescued the low *WNT7A* mRNA expression in DS cells, increasing it to euploid levels, suggesting a potential causal relationship between altered Wnt signaling, neurogenesis, and the over-expression of DYRK1A protein.

The M50 and M54 modules exhibited higher T21 expression, relative to D21, and decreased T21 expression with harmine treatment (Fig. 4C). Consistently, expression of mRNAs in these two modules was elevated in multiple DS brain regions (60). Both modules contain genes with previously annotated functions related to immune response that appear to be dysregulated in DS (117). Four interferon receptor subunits, IFNAR1, IFNAR2, IFNGR2, and IL10RB, are encoded on HSA21, and have been shown to be upregulated by ~1.5-fold on the protein level in DS cells (118). Upregulated interferon receptor expression in DS is strongly implicated in the hyperactive response to interferon stimulation found in skin fibroblasts, lymphoblastoid lines, primary monocytes, and T cells (118). A recent large-cohort proteomics study demonstrated that DS individuals exhibit higher circulating levels of pro-inflammatory cytokines (119).

Altogether, these results reveal that T21 dysregulated protein expression in COs maps to specific human DS brain gene co-expression patterns that are discordant between DS and euploid individuals, particularly during prenatal developmental stages. Moreover, a number of these T21-discordant modules are enriched in proteins with functions related to neurogenesis, neuronal differentiation, and immune response, processes that are known to be disrupted in DS. Proteins within these modules respond to harmine treatment, leading to normalization of expression levels toward D21 levels.

### Harmine upregulated proteins are enriched in a diverse array of functions, cell type-specific markers and transcription factor targets

We next performed a large-scale enrichment analysis on differentially expressed proteins across all stages using the EnrichR libraries (Experimental Procedures). In agreement with the module analysis, we found that harmine upregulated proteins in T21 COs are enriched in markers of neurogenic precursors found in human fetal brain, and markers associated with human cortical brain structures (supplemental Fig. 6B, 7A). Harmine-upregulated proteins were also highly enriched in markers of neuronal subcellular structures, functions related to neuronal differentiation and migration, and regulatory targets of transcription factors and chromatin remodelers with crucial roles in forebrain development (DPF1, MYT1, POU3F3, TBR1, REST, NFATC4) (Column CO_T21_HarmineUP, supplemental Fig. 7A). Taken together, this analysis suggests that harmine treatment leads to large-scale activation of gene expression programs in T21 COs associated with neurogenesis and neuronal differentiation (supplemental Fig. 7B).

### Harmine modulated proteins are enriched in complexes and pathways involved in chromatin remodeling, WNT signaling, and neurogenesis

To visualize functionally-related differentially expressed proteins that may participate in protein complexes or regulatory interactions, we mapped the harmine-modulated differentially expressed proteins (q < 0.1) found in T21 COs onto a STRING interaction network that combines experimentally verified physical interactions with functional interactions found in the literature (81). We then clustered the resulting network to identify densely connected and functionally related subnetworks, and visualized the expression ratios for the T21 vs. D21 effect (node border color) and the T21+harmine effect (node color) (Fig. 5A). One of the larger subnetworks (Fig. 5B) contained proteins enriched in functions related to neurogenesis (q-value = 7 × 10^−12^) and WNT signaling (q-value = 3 × 10^−10^) (for enrichment results of all clusters, see supplemental Data File 1). A number of these proteins are subunits of multiprotein complexes involved in chromatin remodeling. For example, we found four subunits of the NuRD complex (CHD3, HDAC2, MBD3, GATAD2B), which is required for layer specification in the developing cortex (Fig. 5B)(115). Nodes for these subunits were connected to differentially expressed proteins NOTCH1, GLI3, BCL11B and TCF7L2, suggesting important functional relationships between chromatin remodeling and Notch, sonic hedgehog and Wnt signaling in the harmine response of T21 COs. The number of proteins in the two subnetworks in Figure 5B that showed a “harmine normalization” effect was more than 2-fold greater than non-normalized (95 vs. 45 out of 139 proteins total). While harmine leads to only a partial normalization of T21-discordant expression, our evidence suggests that harmine inhibition of a critical target, potentially DYRK1A, results in widespread normalization of protein abundance in T21 COs, particularly for central regulatory transcription factors and morphogens involved in neurogenesis and neuronal differentiation.

## DISCUSSION

We describe a high-depth, quantitative proteomics study of human COs grown from trisomy 21 and euploid iPSC lines, in which we evaluate the proteome-wide effects of a DYRK1A inhibitor, harmine, at four stages of development, starting with iPSC and culminating at COs. On a phenotypic level, we observed a decreased mitotic index phenotype in the T21 iPSC line that agrees with the previous characterization of the C2-4 line (63). Reduced proliferative capacity in human DS lines has been documented previously, notably for iPSC-derived neural precursor cells (82) and primary fibroblasts (120). Consistently, we found that this phenotype could be reversed by long-term treatment with sub-micromolar doses of the DYRK1A inhibitor harmine.

The proteome depth of our organoid trajectory is comparable to a recent comprehensive brain-region resolved proteomics study of mouse brain that identified 13,061 proteins across all regions and cell types (121). We found that the dominant variation in protein expression over the developmental trajectory occurred during the NS to CO transition, when neuroepithelial cells undergo asymmetric neurogenic divisions to form radial glia and early born neurons(94), a process that has been modeled in COs (122). By comparing proteome profiles from euploid COs to T21 COs grown with and without 300nM harmine, we identified a number of cell fate-determining pathways and functional gene signatures that were dysregulated in T21 COs, and normalized by harmine treatment. These include proteins involved in chromatin remodeling and alternative splicing, both of which play critical roles in regulating neurogenesis and forebrain development (123). Most striking was the large number of differentially expressed proteins upon harmine treatment in the T21 COs compared to the earlier iPSC, EB, and NS stages, particularly for proteins associated with early and late differentiated neurons, synaptogenesis, and ECM proteins with known roles in neuronal migration. This stark difference in molecular responses to harmine in the different stages point to the crucial importance of cellular and developmental context, and highlights a distinct advantage of modeling human neurodevelopmental disease states in COs for evaluating potential therapeutic interventions.

The proteome-wide effects, as revealed by high-depth proteomic profiling, combined with the basic cellular phenotypes impacted by harmine treatment, support the view that CO models can provide valuable insights into the molecular mechanisms at work in complex developmental disorders, and suggest critical nodes to target for further investigation. We acknowledge that our analyses examine a single set of tissues derived from two unrelated individuals. Thus, we cannot rule out that a subset of differentially expressed proteins in the T21 vs. D21 comparison are due to genetic background differences. To mitigate this problem, we integrated our proteomics data with a large-scale developmental transcriptome in human DS brain tissue (60), which enabled us to identify those proteomic changes that are recapitulated and discordant with *in vivo* expression patterns. This approach provided us with a computational integrative framework to identify potentially important mechanisms in the altered cortical development of DS that could serve as potential targets for therapeutic interventions. Many differentially expressed proteins observed in our CO model map to discordant gene expression modules in human fetal DS brain. Among the high-value targets in our analysis are those involved in morphogen signaling, neurogenesis, and the specification of laminar identities of neurons in the developing forebrain. The general trend from our data is that the T21-associated changes in protein expression for key regulators of neurogenesis can be partially normalized by harmine treatment. This result corroborates a previous study showing that shRNA knockdown of DYRK1A to euploid levels in a DS iPSC neural induction model rescued impaired neurogenesis and low *WNT7A/B* mRNA expression (82).

Consistently, many proteins in the Wnt, Notch and Sonic Hedgehog (Shh) pathways were differentially expressed in T21 COs and responded to harmine treatment. Hedgehog signaling is attenuated in a number of mouse and human cell DS models, and Shh pathway agonists have been shown to rescue these deficits (124-126). The Notch and Wnt pathways have also been implicated in DS neuropathologies (124-126). DYRK1A has been shown to modulate Wnt (127), Notch (128) and Shh (128) directly, implicating DYRK1A in mediating the effects of harmine documented here. As monoamine oxidase A and DYRK1B activities are also inhibited by harmine at sub-micromolar concentrations (85, 129), we cannot rule out that effects reported here are the result of off-target inhibition. Ongoing efforts are focused on corroborating the effects of harmine using more selective and structurally diverse small molecule inhibitors of DYRK1A, combined with genetic approaches to ablate DYRK1A expression. While DYRK1A is only one of many genes on HSA21 that have been implicated in DS phenotypes (120, 130-133), a large body of research has implicated altered DYRK1A dosage in driving at least some of the key aspects of atypical neurodevelopment in DS, making it an ideal candidate to evaluate in a human stem cell derived CO model of DS brain development.

Collectively, the quantitative changes in T21-specific signaling pathway components and molecular marker profiles, as well as the effects of harmine treatment on the proteome level, are consistent with a complex spectrum of dysregulation in neurogenesis, differentiation, chromatin structure and axon guidance. Our results also point to T21-altered morphogen signaling and ECM remodelling that may play a key role in the dysregulated production of cortical neurons and altered network connectivity observed in Down syndrome, highlighting the utility of deep proteome profiling to quantify molecular aspects of drug efficacy in a human aneuploidy disease model.

## ACKNOWLEDGEMENTS

This work was supported by the Linda Crnic Institute for Down syndrome and a cooperative agreement from DARPA (W911NF-14-2-0019). We thank David W. Russell (UW) for generation of the C2-4 iPSC cell line, University of Colorado Cancer Center Molecular Pathology Shared Resource (CCSG NCI P30CA046934) for karyotyping analysis, Tom Blumenthal and Zachary Poss for insightful discussions, Dr. James Orth and the MCDB microscopy core, Jianli Shi for help on early flow cytometry studies, R. Burgundy for helpful discussions, and Maria Pagratis for moral support.

---

**Supplemental Figure 1:**
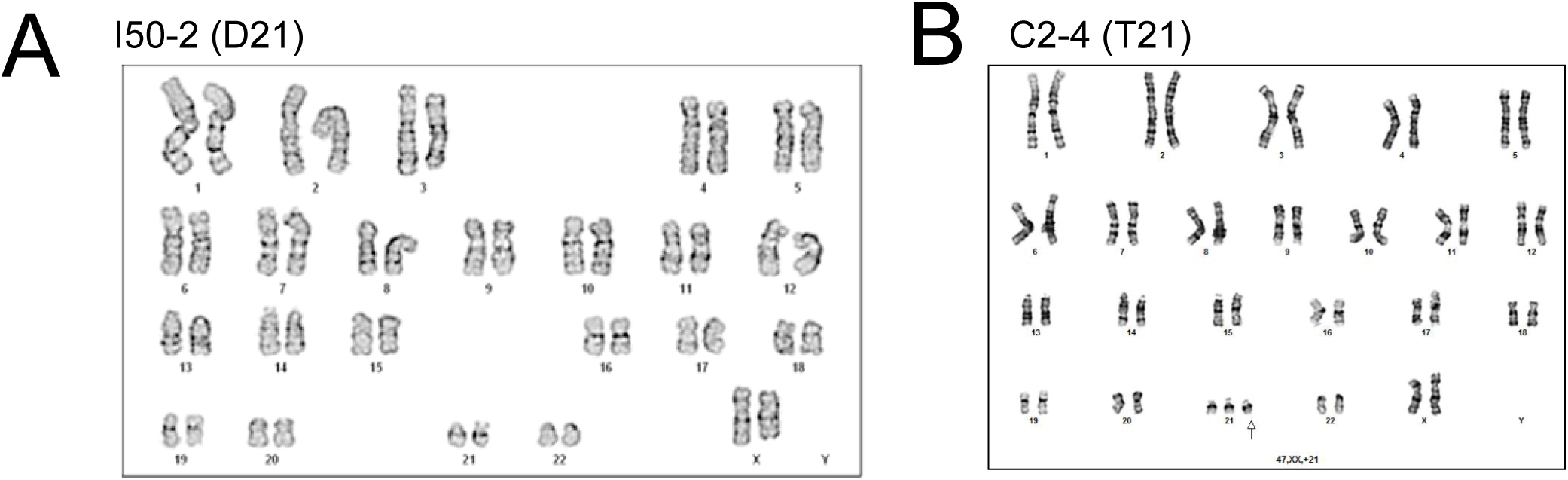
Characterization of iPSC I50-2 (D21) and C2-4 (T21) lines. Karyotypes for iPSC lines. **(A)** I50-2 (D21) exhibited normal female karyotype. **(B)** C2-4 (T21) exhibited trisomy 21 female karyotype.

**Supplemental Figure 2:**
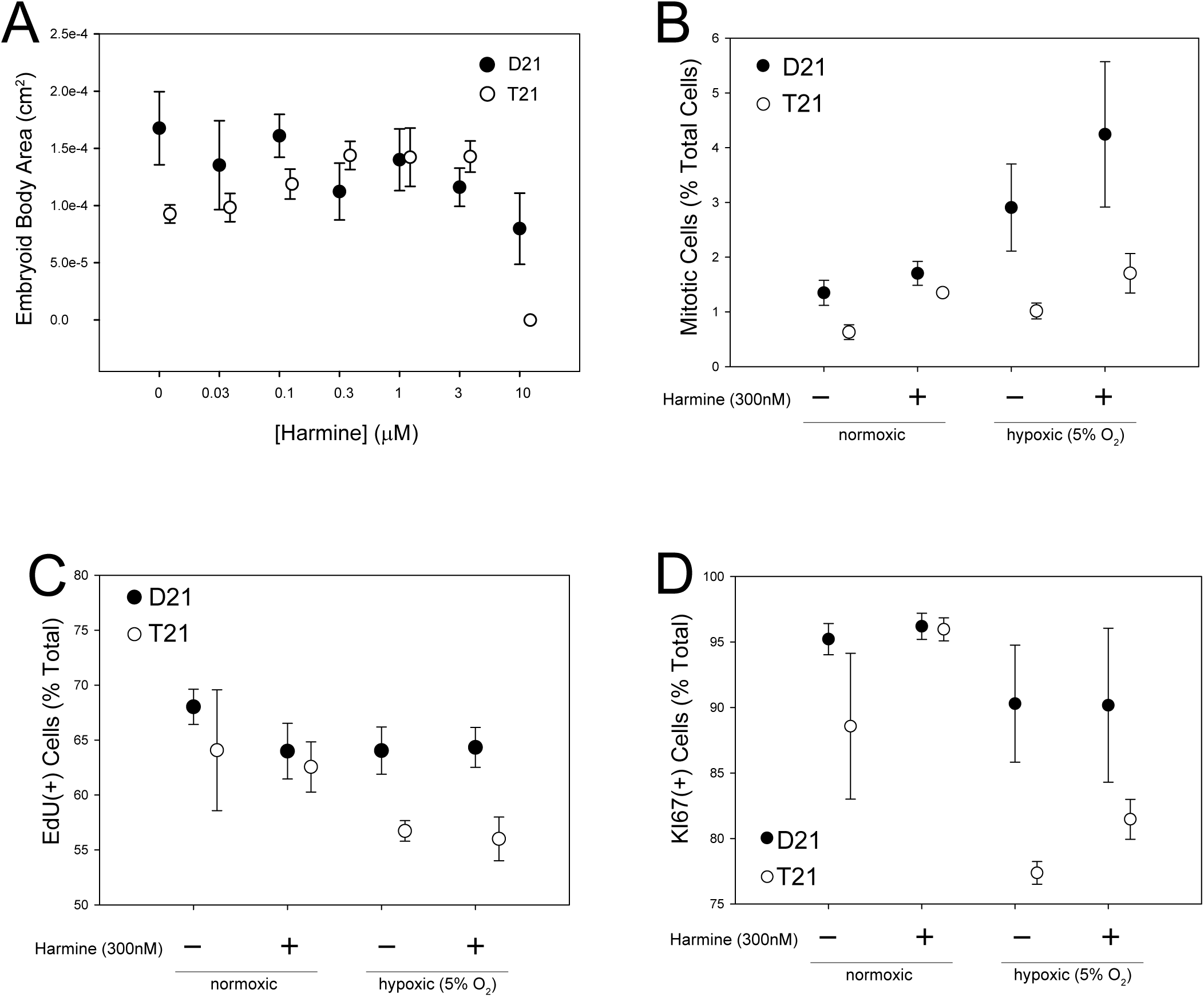
Dose-dependent harmine rescue of T21 embryoid body size distributions and iPSC mitotic index. (**A)** Treatment of iPSC in suspension culture for EB formation with 0-30 μM harmine shows a dose-dependent effect on the calculated area (ANOVA, Main Effect: Harmine F(5, 558)=2.36, p=0.039) of EBs formed from T21 (C2-4) iPSCs, whereas no significant effect of harmine was observed in the D21 line (euploid I50-2) up to 3μM harmine (ANOVA, Main Effect: Harmine F(6,380)=1.15, p=0.33). Harmine concentrations above 10μM were toxic to both lines. **(B)** Treatment of iPSC with 300nM harmine normalized the mitotic index of the T21 cell line (C2-4) with no significant effect on the D21 line (euploid I50-2) in either atmospheric or 5% O2 (3-way ANOVA; Main Effect: Harmine, F(1,21)=12.86, p=0.003; Harmine x Hypoxia Interaction: F(1,21)=0.13, p=0.73). **(C)** There was a significant T21 effect on the percentage of cells that stained positive for EdU (3-way ANOVA; Main Effect: T21, F(1,21)= 6.58, p=0.022), but no significant effect of harmine or a T21 x Harmine interaction on EdU incorporation was observed. **(D)** Harmine had no effect on the ratio of Ki67-positive cells under atmospheric oxygen culture conditions (3-way ANOVA; Main Effect: Harmine: F(1,21)=2.14, p=0.17); similar to the effects observed for EdU+ cells under hypoxic conditions, the T21 iPSC are discordant with respect to the euploid cells in the ratio of Ki67-positive cells (3-way ANOVA; Main Effect: T21; F(1,21)=11.36, p=0.005).

**Supplemental Figure 3:**
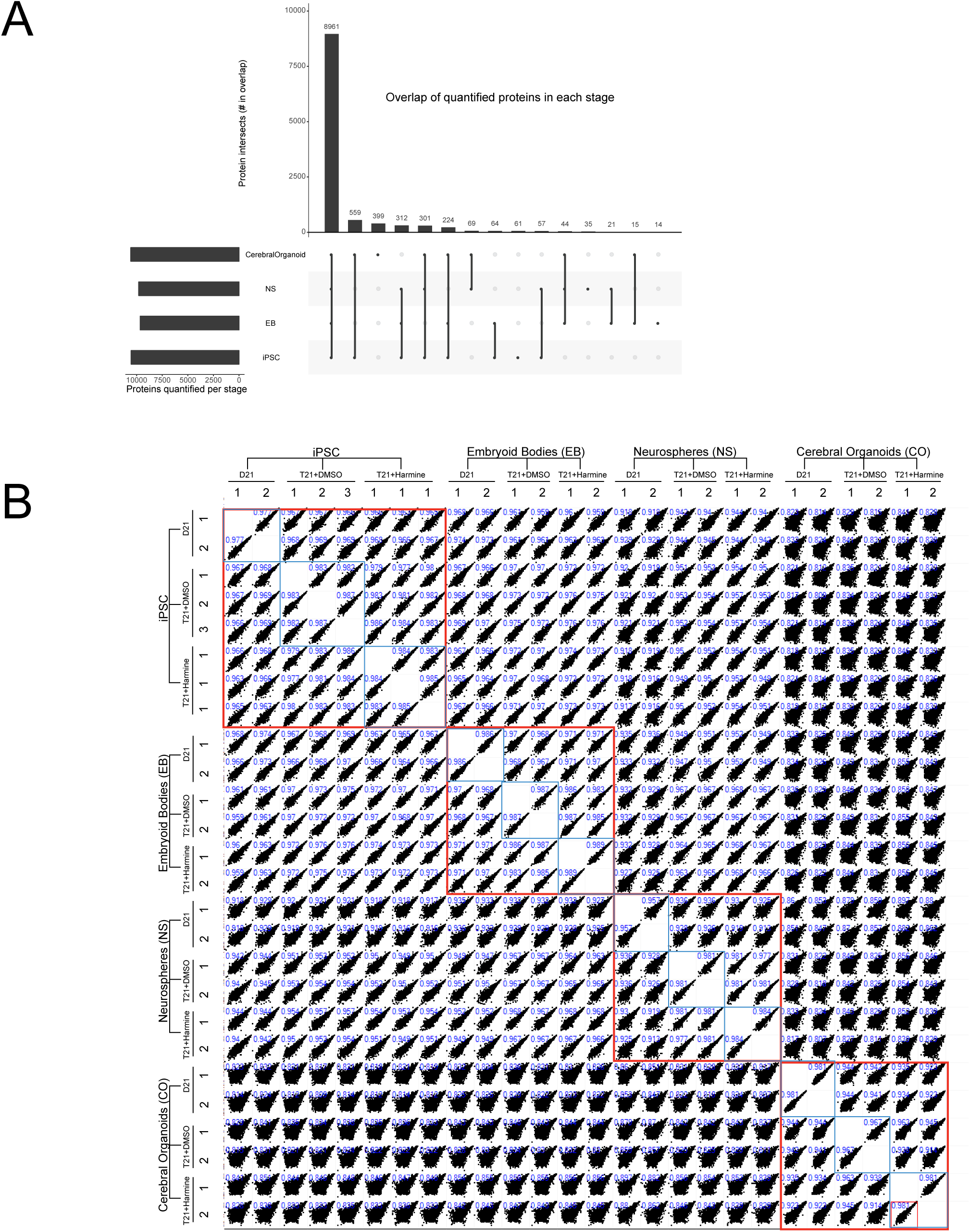
Overlap of quantified proteins across four developmental stages of iPSC differentiation. **(A)** UpsetR plot, which shows the protein groups, represented as uniprot identifiers, quantified in at least three samples that overlap between different stages. UpsetR plots are scalable alternatives to Venn diagrams (Conway, J.R. et al. 2017). A Protein group was considered quantified for this analysis if a valid LFQ value was found at least three samples across all 26 samples. Lines connecting black dots indicate overlap, and the bar indicates the magnitude of the overlap. **(B)** Scatter plot matrix of protein group LFQ values showing that within stage and within biological replicate group variability is less that between stage variability. Biological replicate numbers are shown below the annotations for cell line, treatment condition, and stage. Pearson correlation coefficient for each pairwise sample comparison is shown in blue in each panel.

**Supplemental Figure 4:**
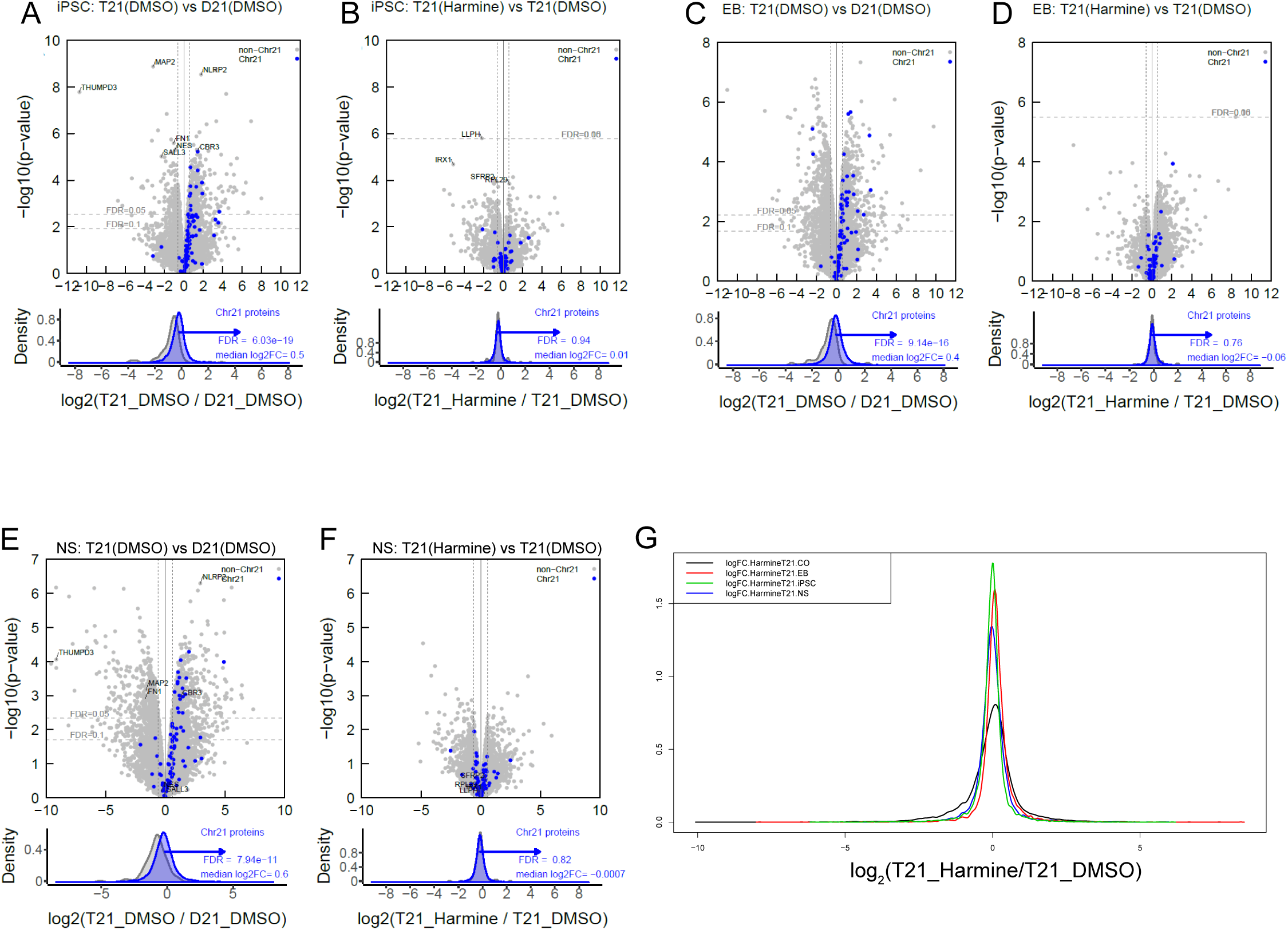
Differential expression for T21 vs. D21 and T21+Harmine effects in each of four developmental stages. Shown in A-F are volcano plots for the T21+DMSO vs. D21+DMSO comparisons (A, C, E) at the iPSC stage (A,B), embryoid body (EB) stage (C,D) and neurosphere (NS) stage (E,F).

**Supplemental Figure 5:**
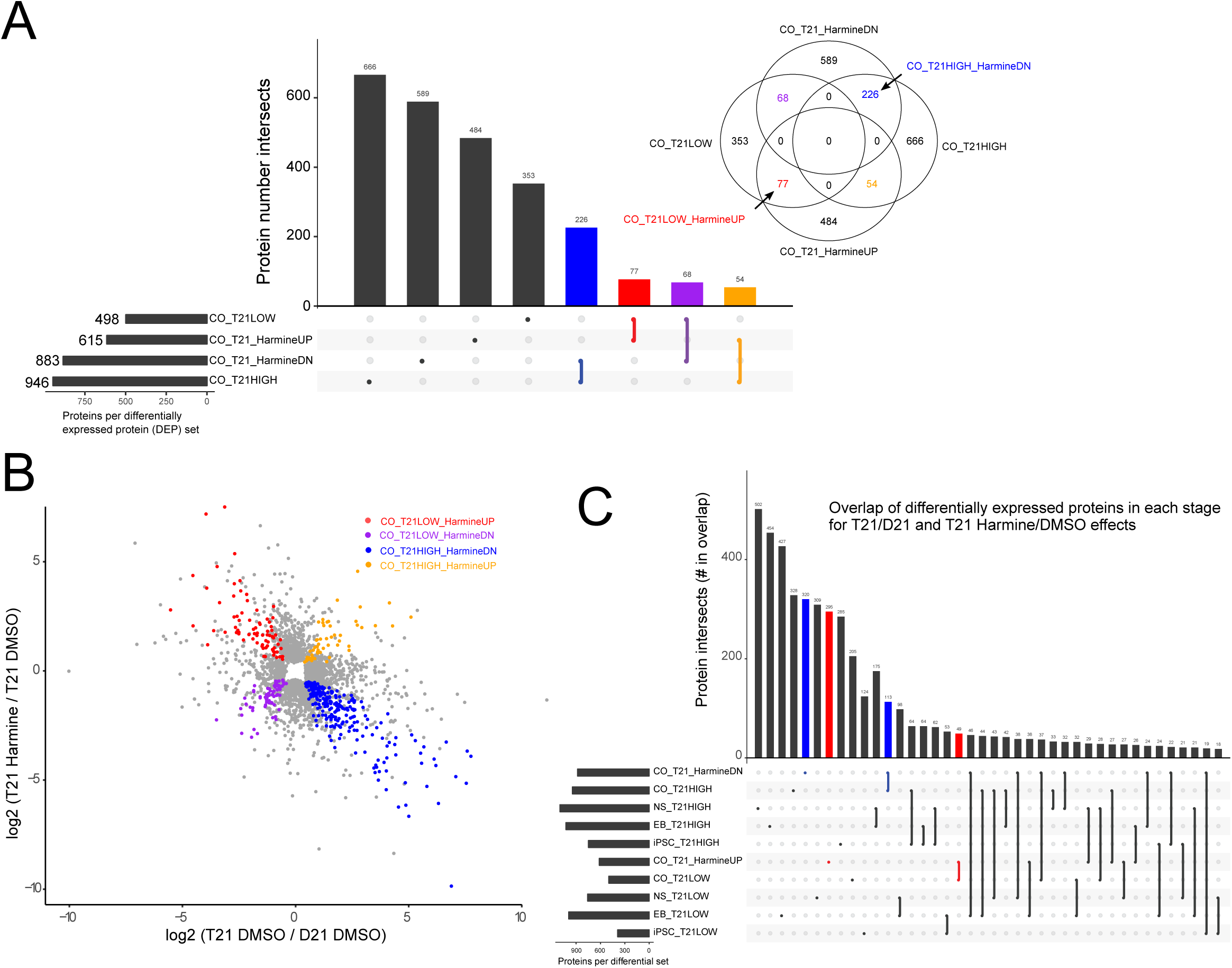
**Overlap of differentially expressed protein (DEP) sets,** for the (**A**) organoid stage and whole trajectory data set. (**B**) Scatter plot of the two differential expression comparisons, as in Fig. 4A, showing differentially expressed proteins within the four binary intersections (q < 0.1) indicated by color in the bar chart and Venn diagram in (A). In (**C**), red and blue indicate DEP sets and interactions discussed in the main text with harmine upregulated and harmine downregulated proteins, respectively.

**Supplemental Figure 6.**
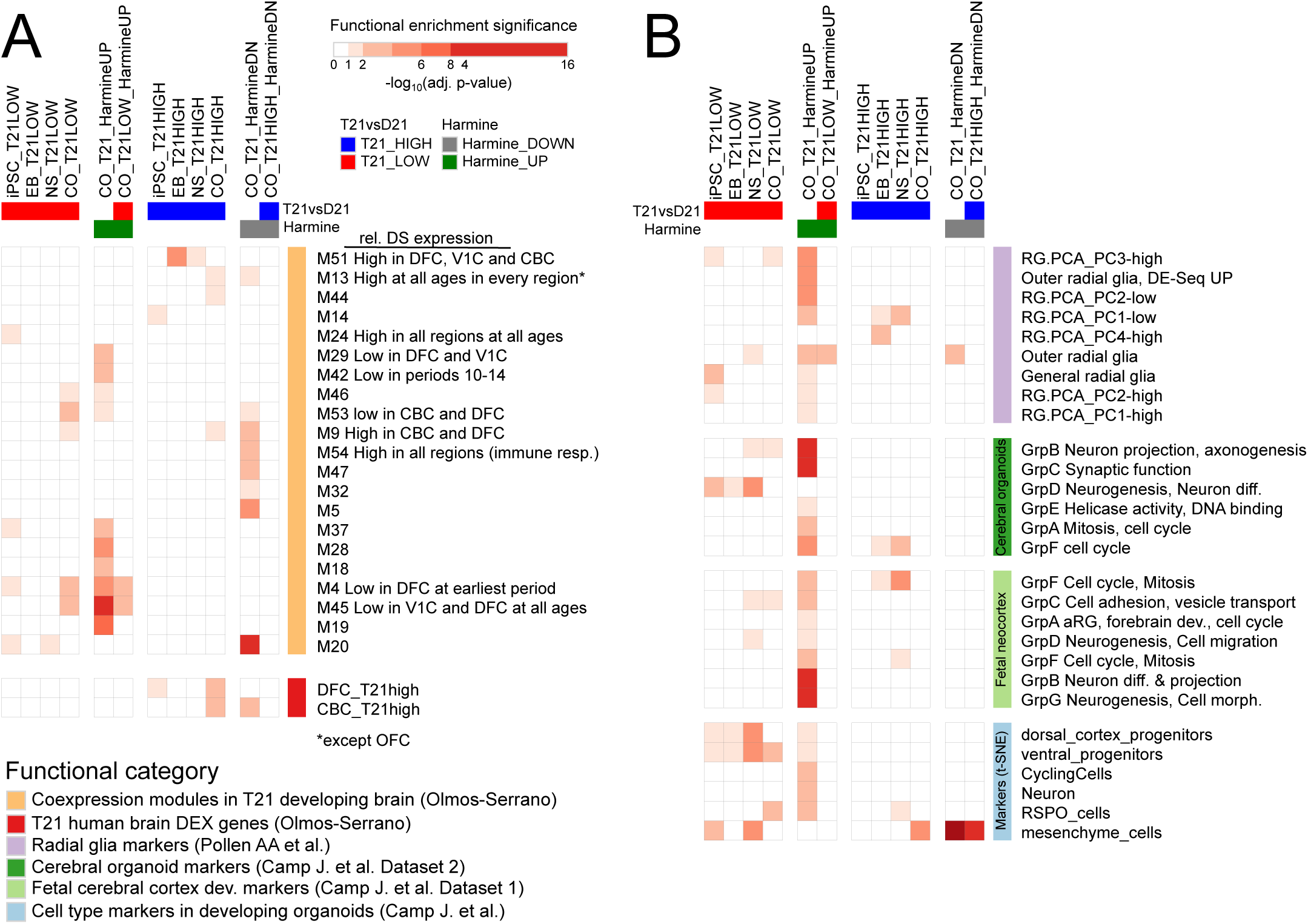
Trisomy 21 differentially expressed proteins and harmine-modulated proteins map to expression signatures identified in transcriptome studies of human forebrain development. Functional enrichment analysis of differentially expressed proteins in T21 vs. D21 at each developmental stage and harmine-treated T21 cerebral organoids, using the Fisher exact test, with color indicating the significance level. Gene set enrichment analysis was performed for the proteins showing differential expression in the T21 vs. D21 and T21 harmine comparisons (q < 0.1), with total numbers in each differentially expressed protein category as reported in Table 1. P-values resulting from the enrichments in all categories shown were collectively Benjamini-Hochberg corrected for each differential category (column). (**A**) Functional enrichment of differentially expressed proteins against region-specific differentially expressed genes identified in Olmos-Serrano et al. (2016) and associated functional modules from the same study. (**B**) Enrichments for cell type specific gene sets. oRG, outer radial glia, vRG, ventral radial glia. The nomenclature for DEP sets used in enrichment analysis are as follows, with q < 0.10 for all sets: *stage*_T21HIGH refers to the differentially expressed proteins showing higher expression in the T21 vs. D21 comparison for that specific stage. Conversely, *stage*_T21LOW refers to the differentially expressed proteins showing lower expression in the T21 vs. D21 comparison for that specific stage. CO_T21_HarmineUP refers to differentially expressed proteins showing a significant increase in the T21 COs upon harmine treatment. CO_T21_HarmineDN refers to differentially expressed proteins showing a significant decrease in the T21 COs upon harmine treatment. CO_T21LOW_HarmineUP is the intersection of CO_T21LOW and CO_T21_HarmineUP, and CO_T21HIGH_HarmineDN is the intersection of CO_T21HIGH and CO_T21_HarmineDN. The full set of terms and statistics can be found in Supplemental Table 4.

**Supplemental Figure 7.**
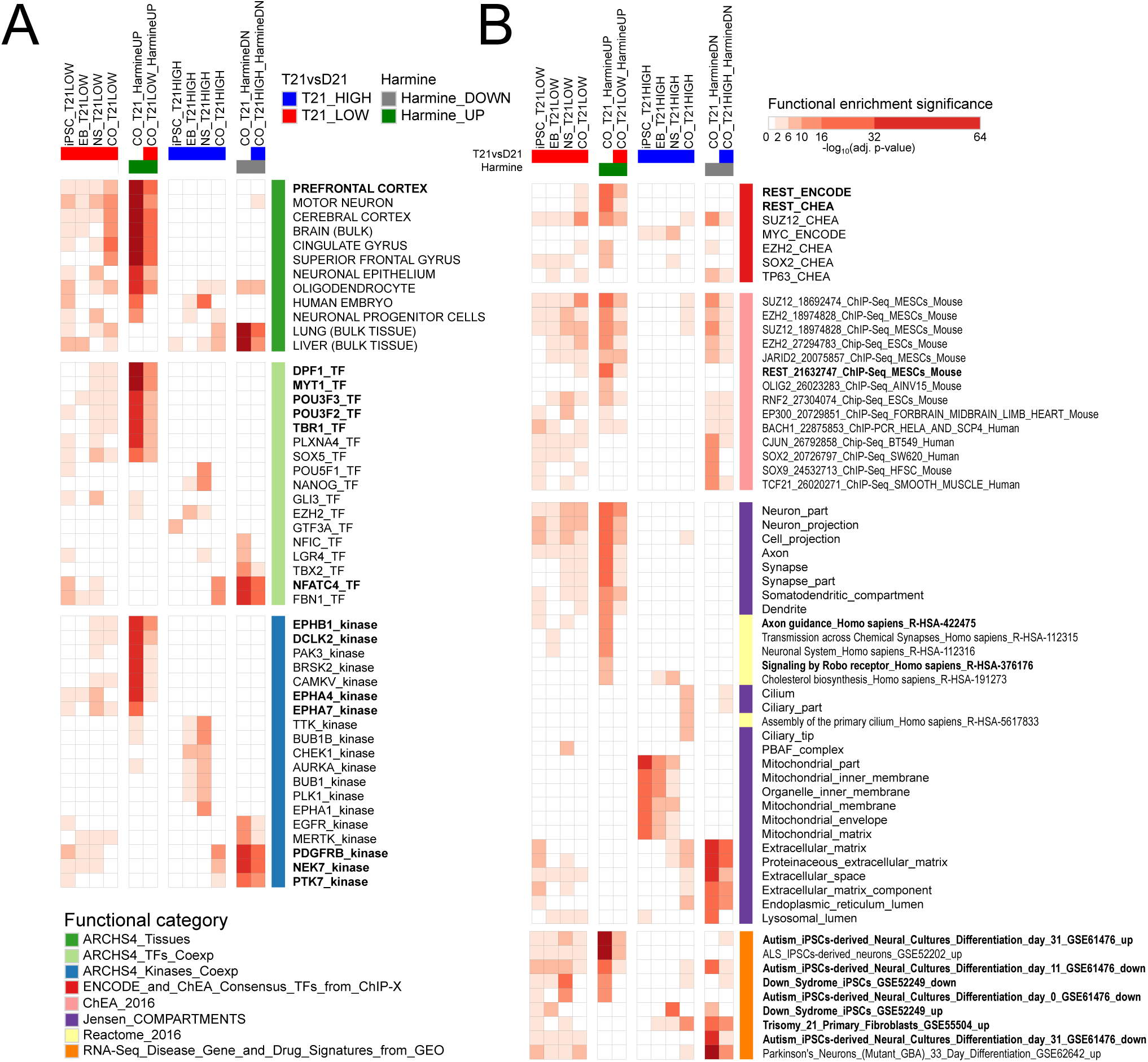
Trisomy 21 and harmine-treatment differential proteins map to transcription factor and kinase co-expression modules and transcriptional repressor targets. (**A**) Functional enrichment of DEP sets described in Fig. 5 against gene sets in the ARCHS4 compendium. (**B**) Enrichment analysis against GEO differential expression data sets, transcription factor binding targets, and ChIP-seq data sets using libraries compiled for the EnrichR tool (http://amp.pharm.mssm.edu/Enrichr/#stats). To identify signatures of disease related processes that overlap with differentially expressed proteins, we tested for enrichment in disease gene signatures using a compendium of disease associated differential gene sets from GEO data sets that have been partitioned into up-regulated and down-regulated gene sets with respect to the disease state (Lachmann et al. 2018). The GEO data sets shown were selected for significance and disease relevance from a larger set of statistically significant enriched terms (the full set of enrichments and statistics can be found in **Supplemental Table 5**).

## ABBREVIATIONS

HSA21: human chromosome 21
DS: Down syndrome
CO: cerebral organoid
EB: embryoid body
iPSC: induced pluripotent stem cell
DMSO: dimethyl sulfoxide
EdU: 5-ethynyl-2’-deoxyuridine
FDR: false discovery rate
T21: the trisomy 21 C2-4 iPSC line
D21: the euploid I50-2 iPSC line
TX100: Triton X-100; NS, neurosphere
DPBS: Dulbecco’s phosphate-buffered saline
PBS: phosphate-buffered saline
AGC: Automatic Gain Control
LFQ: label-free quantitation
PCA: principal component analysis
ECM: extracellular matrix
WGCNA: weighted gene co-expression network analysis

